# Dysregulated lymphocyte localization in idiopathic multicentric Castleman disease

**DOI:** 10.64898/2025.12.12.693846

**Authors:** Melanie D. Mumau, Michael V. Gonzalez, Katherine S. Forsyth, M. Betina Pampena, Abiola H. Irvine, Criswell L.M. Lavery, Joseph M. Zinski, Ira D. Miller, Sam Barnett Dubensky, Derek A. Oldridge, Laura A. Vella, Michael R. Betts, David C. Fajgenbaum

## Abstract

Idiopathic multicentric Castleman disease (iMCD) is a rare and life-threatening hematologic disease involving episodic flares of uncontrollable systemic inflammation by unknown causes. Hallmark features of iMCD include multiple enlarged lymph nodes with characteristic histopathological phenotypes and a potentially fatal, cytokine release syndrome. The key pathophysiologic drivers of disease are poorly understood and few effective treatment options exist. Here, we discovered an association between elevated chemokines, namely CXCL13, and lymph node size in iMCD, providing one possible explanation for the lymphadenopathy observed clinically. Instead of a concurrent increase in circulating CXCL13 and CXCR5-expressing cells that has been described in other contexts, during active disease, chemokine-responsive lymphocytes downregulated CXCR5 levels in iMCD. Despite heightened chemokine production by lymph node stromal cells, T and B cells failed to appropriately respond to their cues locally within the tissue and were particularly scarce within CXCL13-expressing germinal centers (GC). Inflammatory signals in iMCD lymph nodes appeared to restrict the production and movement of T follicular helper cells, which play an important role in facilitating appropriate GC responses. Together, these data provide a link between dysregulated chemokine production and germinal center lymphocyte trafficking, highlighting a potential mechanism and therapeutic target in iMCD lymphadenopathy.

**One Sentence Summary:** Lymphocyte chemotaxis to discrete areas of lymphoid tissue is disrupted in idiopathic multicentric Castleman disease.

## INTRODUCTION

The last decade has brought major advancements in the understanding, diagnosis, and treatment of iMCD (1–3). Diagnostic criteria for iMCD requires multiple enlarged lymph nodes with a range of histopathologic features that include atrophic or hyperplastic germinal centers (GCs), follicular dendritic cell (FDC) prominence, plasmacytosis, and vascularity (1). However, there is a limited understanding of what causes the observed lymphadenopathy and its associated histopathological features, as well as their mechanistic implications. There are three known iMCD subtypes, which range in severity, clinical presentation, and histopathology. The least severe subtype of iMCD, idiopathic plasmacytic lymphadenopathy (iMCD-IPL), is associated with thrombocytosis, hypergammaglobulinemia, and lymph node plasmacytosis, with patients experiencing moderate symptomatology (4, 5). The most life-threatening iMCD subtype presents with acute and severe symptoms, including thrombocytopenia (T), anasarca (A), fever (F), reticulin fibrosis or renal failure (R), and organomegaly (O), collectively known as iMCD-TAFRO (5–7). Lymph node histopathology of iMCD-TAFRO patients typically involves hypervascularity with regressed GCs and FDC prominence. The third subtype, iMCD-not otherwise specified (iMCD-NOS), is reserved for patients who meet the iMCD diagnostic criteria but do not satisfy the requirements for the other two subtypes.

Despite these clinical classifications, little is understood about iMCD etiology and mechanisms, i.e. the key dysregulated signaling pathways, cell types, and immune mediators that drive disease pathogenesis and the life-threatening cytokine storm associated with iMCD-TAFRO patients. While germline mutations, somatic mutations, as-yet-identified antigens, and auto-antibodies have all been hypothesized to potentially cause iMCD, neither shared genetic linkages or associated pathogens have been discovered (8). A recent study discovered the presence of common connective tissue autoantibodies in iMCD, but no specific autoantibodies were identified (9). Mechanistically, IL-6 and mTOR signaling have been shown to be important mediators of iMCD-TAFRO, as a portion of patients respond to treatment with IL-6 or mTOR inhibitors (3, 10). However, ∼50% of iMCD-TAFRO patients do not achieve long-term remission with these types of treatments, suggesting gaps in our understanding of iMCD-TAFRO with important implications for patient outcomes (3). Treatment-refractory patients may initially respond to IL-6 blockade or other therapies and enter a brief remission period to then relapse a short time thereafter (10, 11). More treatment strategies are needed for these patients who fail to achieve long-lasting remission. A better characterization of the immune mediators including cytokines, their involvement in disease pathogenesis, and how they affect key cell types will help to improve the mechanistic understanding of the disease and may uncover new treatment approaches.

Serum proteomic profiling of the blood of iMCD patients previously identified CXCL13 and CCL21 among the top-upregulated proteins in circulation (12–14). However, these studies predominantly utilized iMCD-IPL and iMCD-NOS patients and included only a small sample size of iMCD-TAFRO patients (10/88 patients (12) and 2/6 patients (14)). More studies are needed to evaluate the serum proteome in iMCD-TAFRO. In studies to date, we and others have demonstrated elevated CXCL13 levels by non-hematopoietic/stromal cells in the GC in iMCD (12, 15). Stromal cells in secondary lymphoid organs (SLO) produce CXCL12, CCL19, and CCL21, which along with CXCL13, promote the homing of lymphocytes from circulation into the SLO and to specific regions within them. Specialized stromal cell subsets produce different chemoattractants with fibroblastic reticular cells (FRC) producing CCL19, CCL21 and CXCL12, which attract lymphocytes out of circulation and into the SLO paracortex region. CCL19 and CCL21 also act locally within the tissue, where CCR7-expressing cells follow their concentration gradients, with the highest levels typically expressed in the interfollicular (IF) space. High concentrations of CXCL13 by FDC attract CXCR5-expressing cells to B follicles at steady state and direct them towards the light zone region of GCs during inflammation. As multiple enlarged lymph nodes and dysmorphic histopathology are hallmarks of iMCD, we hypothesized that high levels of chemoattractants in iMCD contribute to the lymphadenopathy observed and affect the cell types that respond to these signals, such as T and B lymphocytes.

In this study, we demonstrate that during active iMCD, chemokines crucial for trafficking to and within lymph nodes were elevated in the serum and lymph nodes of iMCD-TAFRO patients and may be responsible for the associated lymphadenopathy and histopathological changes. Flow cytometry, gene expression data, and immunofluorescence studies revealed that, despite high levels of chemokines in iMCD-TAFRO, lymphocyte localization within the lymph node was dysregulated. We discovered that enhanced type I IFN signaling may restrict the ability of T cells to appropriately localize within GCs and limit Tfh differentiation. The data presented here provide insights into potential mechanisms underlying iMCD-TAFRO, the most severe and life-threatening subtype of iMCD.

## RESULTS

To better understand the mechanisms that promote pathogenesis in iMCD-TAFRO, we performed serum proteomic analysis on a cohort of iMCD-TAFRO patients with active disease (n = 24) and healthy donors (n = 15). We quantified 6,408 analytes and discovered that factors promoting chemotaxis to and within SLO were among the most up-regulated proteins in iMCD-TAFRO compared to healthy donors. CXCL13 was the second most significantly up-regulated protein profiled, more than 7-fold higher (log2 fold-change = 2.92) in iMCD-TAFRO than healthy controls, and CCL21 and CCL19 were among the top 20 most up-regulated cytokines and chemokines (Figure 1A-F). Although not as pronounced, CXCL12 was also increased in iMCD-TAFRO (Figure 1C). Utilizing a previously published serum proteomic dataset to determine if these findings were non-specific inflammatory changes (13), we compared levels of CXCL12, CXCL13, CCL19, and CCL21 between iMCD-TAFRO (n = 11) and related lymphadenopathies that included HHV8^+^ MCD (n=20), lymphoma (n=20), and rheumatoid arthritis (RA) (n = 19) (Figure 1G-I and Supplemental Figure 1A). All 4 chemokines were significantly elevated in iMCD-TAFRO compared to RA and HHV8^+^ MCD. Interestingly, CCL21 was approximately 2-fold higher compared to all lymphadenopathies tested, including lymphoma. To determine whether the increase in serum chemokines in iMCD-TAFRO contributed to the characteristic lymph node enlargement, we investigated the relationship between lymph node size and the levels of CXCL12, CXCL13, CCL19, or CCL21. We utilized our serum proteomic data and corresponding clinical data from iMCD-TAFRO patients and discovered significant, positive correlations between CXCL13, CCL19, and CCL21 levels, but not CXCL12, and lymph node size measurements (Figure 1J-L and Supplemental Figure 1B).

**Figure 1.**
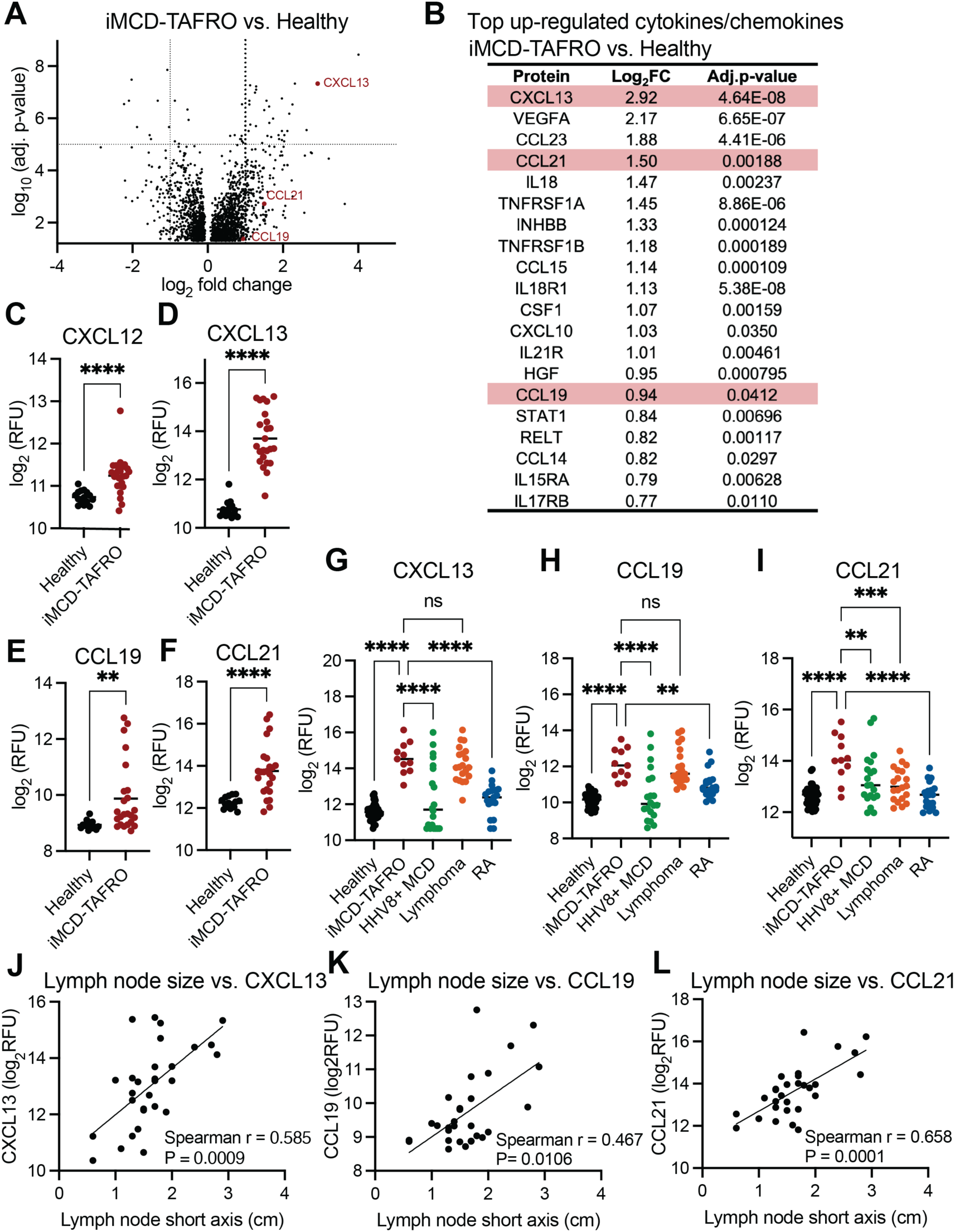
Elevated serum chemokines associated with lymphadenopathy in iMCD-TAFRO. The serum proteome (>7,000 analytes) was profiled in serum samples from patients. (A) Volcano plot comparing all analytes in blood from iMCD-TAFRO patients (n=24) versus healthy controls (n=15). (B) Top 20 up-regulated cytokines and chemokines in iMCD-TAFRO compared to healthy donor controls. Comparison of individual analytes, (C) CXCL12, (D) CXCL13, (E) CCL19, (F) CCL21 in iMCD-TAFRO patient samples versus healthy donor controls. (C) In a second study, (G) CXCL13, (H) CCL19, and (I) CCL21 levels were compared between iMCD-TAFRO and other inflammatory diseases (Healthy, n=42; iMCD-TAFRO, n=11; HHV8+ MCD = human herpesvirus-8 associated MCD, n=20; Lymphoma = Hodgkin’s lymphoma, n=20; RA = rheumatoid arthritis, n=19. Dots in aforementioned graphs represent individual patient samples. Levels of (J) CXCL13, (K) CCL19, and (L) CCL21 were plotted against lymph node sizes taken from clinical radiological data. *p<0.05. **p<0.01 ***p<0.001 ****p<0.0001.

While iMCD has often been referred to as a lymphoproliferative disorder, we investigated whether proliferation also contributed to the characteristic lymphadenopathy in iMCD-TAFRO. We analyzed lymph node size and fluorodeoxyglucose (FDG) uptake from radiology reports and performed immunohistochemistry against Ki-67 on lymph node tissue from iMCD-TAFRO patients (Supplemental Figure 1C-F). Compared to controls that included lymphoma, we discovered lower FDG uptake and Ki-67 staining in iMCD-TAFRO. Furthermore, we did not find an association between lymph node size and proliferation, as assessed by FDG avidity or Ki-67 expression (Supplemental Figure 1G-H). Together, these data suggest that increased trafficking of circulating, chemokine-responsive cells, rather than cellular proliferation, may promote the lymphadenopathy observed in iMCD-TAFRO.

Given that these specific chemokines, CXCL12, CXCL13, CCL19, and CCL21, are known to be secreted from discrete areas of lymph nodes and could promote lymphadenopathy in iMCD by attracting cell types that express cognate receptors, we wanted to investigate a potential source of chemokine production within the lymph node. We utilized available formalin-fixed paraffin-embedded (FFPE) tissue from patients and performed spatial transcriptomics of iMCD-TAFRO lymph nodes and undiagnosed inflammatory or reactive controls. We used the Bruker/Nanostring GeoMx platform to interrogate gene expression in spatially distinct regions—the GC and the interfollicular T cell zone (Figure 2A). Unbiased analysis of the whole transcriptome (>18,000 genes) revealed that CXCL13, CCL19, and CCL21 were among the top up-regulated genes in the GC of iMCD-TAFRO lymph nodes (n=4) compared to controls (n=3) (Figure 2B). Interestingly, CXCL13 and CXCL12 were two of the most upregulated genes in the interfollicular (IF) space, an area where CXCL13 is typically not produced, in iMCD-TAFRO (n = 6) (Figure 2C-D). In the GC, we found that CXCL13 gene expression was approximately 5 times higher (log2 fold-change = 2.3) in iMCD-TAFRO lymph nodes compared to controls (n = 3) (Figure 2E). Gene expression of CXCL13 in the IF space of iMCD-TAFRO lymph nodes was 3.7x higher (log2 fold-change = 1.9) than in controls and, interestingly, was almost as high as CXCL13 expression in the GCs of inflammatory controls. Although CCL19 and CCL21 expression are typically more restricted to the IF space, average expression of CCL21 was 7x (log2 fold-change = 2.8) higher in iMCD-TAFRO GCs compared to controls and although not significant, CCL19 expression was 2.6x higher (log2 fold-change = 1.4) in iMCD-TAFRO (Figure 2E-F). As high levels of CXCL13 are typically expressed by cells within the GC and CCL19, CCL21, and CXCL12 are most notably produced outside of the GC to attract cells out of circulation and into the T cell zone, these data suggest that mechanisms promoting chemotaxis are not only enhanced, but spatially dysregulated with respect to the local pattern of expression within the lymph node in iMCD-TAFRO.

**Figure 2.**
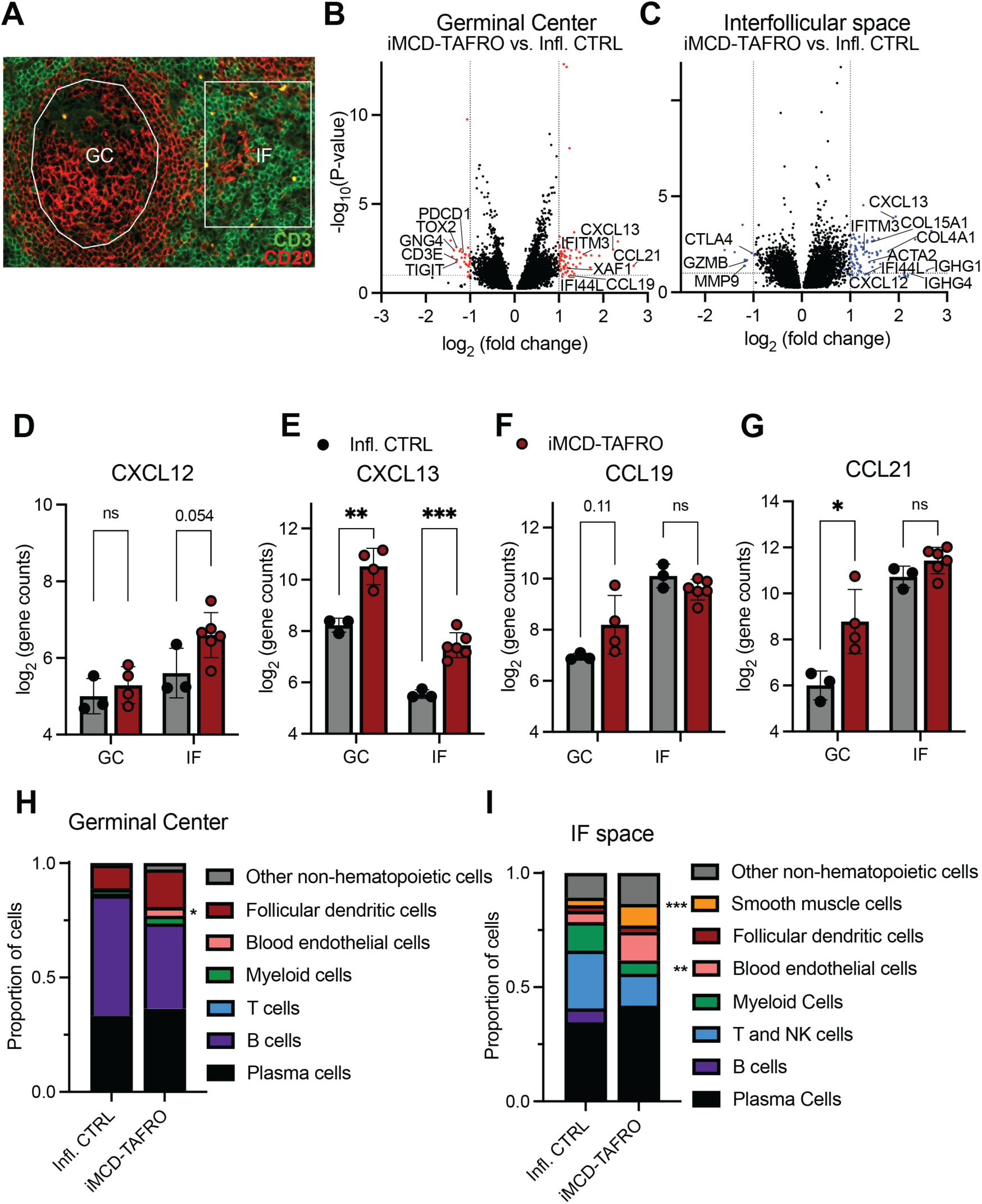
Regional transcriptomics indicates dysregulated expression of chemokines in iMCD-TAFRO lymph nodes. (A) Representative immunofluorescence image taken on the GeoMx spatial profiler to denote specific regions of lymph node tissue (GCs and IF regions) that were analyzed for gene expression. (B) Volcano plots depicting differentially expressed genes within (B) GCs and (C) IF space between iMCD-TAFRO and inflammatory control lymph nodes. Infl. CTRL, n=3; iMCD-TAFRO n=6. Expression of individual chemokines (D) CXCL12, (E) CXCL13, (F) CCL19, and (G) CCL21 in regions profiled. Spatial deconvolution was performed using transcripts collected from (H) GC and (I) IF space regions. Dots in bar graphs represent individual patient samples. *p<0.05. **p<0.01. ***p<0.001.

Although chemokine expression was elevated in iMCD-TAFRO, it was unclear whether differences in cellular heterogeneity contributed to these gene expression differences, as we profiled regions within the tissue, not individual cells. To determine if changes in the cellular composition in the GC and IF regions contributed to the observed increase in chemokine expression in iMCD-TAFRO, we performed spatial deconvolution of the GeoMx datasets for each region in iMCD-TAFRO and inflammatory control lymph nodes. Generally, we observed similar cell frequencies in both regions with some notable differences (Figure 2G-H). We found a significantly higher frequency of FDC in GC of iMCD-TAFRO lymph nodes (Figure 2G), consistent with FDC prominence in iMCD lymph node described by histopathology (1). In the IF space of iMCD-TAFRO lymph nodes, approximately 40% of the cells were predicted to be non-hematopoietic/stromal cells, compared to about 20% in the inflammatory control lymph nodes (Figure 2H). Contributing to these differences were an increase in the frequency of smooth muscle cells and blood endothelial cells (BEC) in iMCD-TAFRO, also consistent with the hypervascular pathology described in iMCD. Together, these data suggest that elevated chemokine expression in iMCD-TAFRO lymph nodes may be caused by an expansion of non-hematopoietic cell types. However, it remains unclear if increased chemokine levels in iMCD-TAFRO are due to population differences or overexpression within individual cells, both of which can be addressed by single-cell RNA sequencing.

Although both hematopoietic and non-hematopoietic cells can express CXCL12, CXCL13, CCL19, and CCL21, we aimed to determine whether the observed increase in their expression was due to changes in cell populations that typically express these chemoattractants and/or caused by an increase in chemokine production on a per-cell basis. To determine the source of elevated chemokine expression in iMCD-TAFRO lymph nodes, we performed single-cell gene expression profiling of gently dissociated FFPE tissue using 10X Genomics’ FLEX pipeline. We utilized lymph nodes from 4 iMCD-TAFRO patients and 4 uncharacterized inflammatory controls and defined major cell types by annotating distinct clusters that included T and B cells, stromal cells, endothelial cells, macrophages, and plasma cells (Figure 3A). To improve the annotation of closely related cell types, we divided our dataset into 4 broad cell categories for finer sub-clustering: non-hematopoietic/stromal cells, T and NK cells, B cells, and myeloid cells. The major producers of CXCL12, CXCL13, CCL19, and CCL21 were non-hematopoietic/stromal populations in both control and iMCD-TAFRO lymph nodes (Figure 3B). Analysis of the non-hematopoietic/stromal sub-cluster revealed distinct populations including FDC, perivascular reticular cells (PRC), and other fibroblast reticular cells (FRC), BEC and lymphatic endothelial cells, and smooth muscle cells (Supplemental Figure 2A-B). Gene expression analysis between non-hematopoietic/stromal cell populations in iMCD-TAFRO and inflammatory control lymph nodes indicated that CXCL12 was a top differentially expressed gene by BEC. CXCL13, CCL19, and CCL21 were among the most up-regulated genes in iMCD-TAFRO FDC (Figure 3C and Supplemental Figure 2C), which was surprising given that FDC typically do not express CCL19 or CCL21. FDC expressed CXCL13 (log2 fold-change = 2.05), CCL19 (log2 fold-change = 2.35), and CCL21 (log2 fold-change = 1.92) approximately 4-times higher in iMCD TAFRO compared to inflammatory controls (Figure 3D). Hematopoietic cell types known to express CXCL13, including T follicular helper (Tfh) cells and macrophages, did not appear to be major contributors to the differences in chemokine expression (Supplemental Figure 2D). Fibroblasts, known to localize to the T cell zone, PRC and FRC, expressed the highest levels of CCL19 and CCL21, which were significantly elevated in iMCD-TAFRO lymph nodes compared to controls. In addition to chemokines, iMCD-TAFRO lymph node stromal cells, notably PRC and BEC, up-regulated *VCAM1* and *ICAM1* (Figure 3E), which are critical for lymphocyte adhesion and extravasation out of circulation and into the tissue. Further examination of the non-hematopoietic/stromal cell subcluster did not reveal any changes in population frequencies, suggesting that chemokine expression is likely up-regulated at the individual cell level (Supplemental Figure 2E).

**Figure 3.**
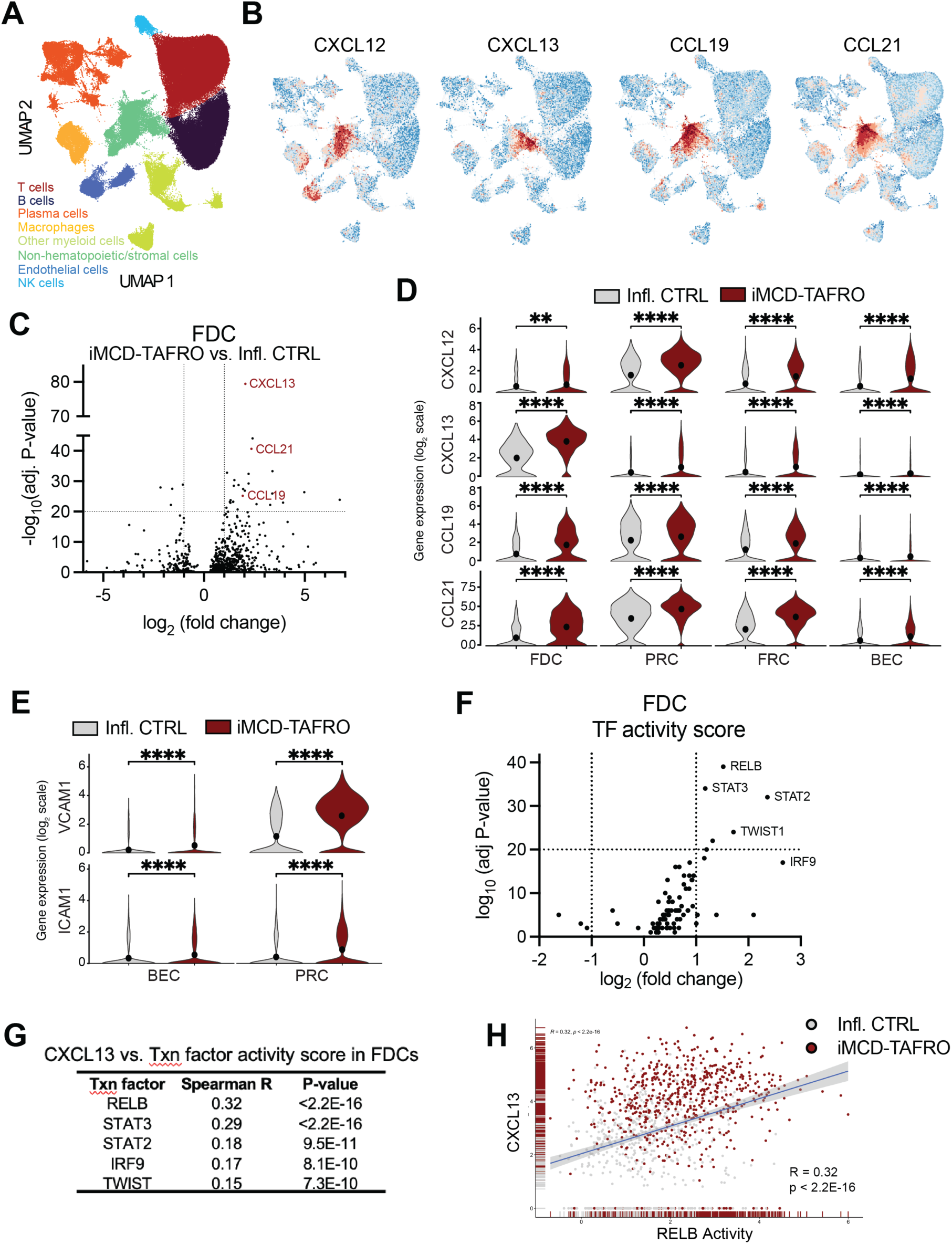
Gene expression profiling of single cells reveals elevated chemokine expression via enhanced inflammatory signaling pathways in iMCD-TAFRO stromal cells. (A) Uniform manifold approximation projection (UMAP) of single-cell RNA sequencing data performed on lymph node cells dissociated from FFPE sections. Infl. CTRL, n=4; iMCD-TAFRO, n=4. (B) Populations were analyzed for gene expression of CXCL12, CXCL13, CCL19, and CCL21. (C) Volcano plot comparing iMCD-TAFRO vs. Infl. CTRL follicular dendritic cells (FDC). (D) Violin plots comparing chemokines in FDC, perivascular reticular cells (PRC), and fibroblastic reticular cells (FRC) in iMCD-TAFRO lymph nodes. (E) Comparisons of expression of adhesion molecules, VCAM1 and ICAM1, in blood endothelial cells (BEC) and PRC. Dots in violin plots represent median values for each gene. (F) Transcription factor activity scores were calculated in iMCD-TAFRO and control FDCs and compared. (G) The most significant transcription factor activity scores were compared to CXCL13 expression in combined FDC from both iMCD-TAFRO and Infl. CTRL. (H) Relationship between CXCL13 and activity of RELB, a target of non-canonical NFkB signaling. **p<0.01. ***p<0.001. ****p<0.001.

To investigate the mechanisms that promote chemokine expression in iMCD-TAFRO, we calculated transcription factor activity scores based on gene expression for FDC, PRC, and FRC (Figure 3F and Supplemental Figure 2F-G). We discovered that several transcription factors, including RELB, TWIST1, STAT3, and IRF9, were predicted to be more active in each iMCD-TAFRO stromal cell population compared to controls. As expected, several of these transcription factors, like RELB and IRF9, are involved in CXCL13 induction (16, 17). To connect transcription factor activity with transcription of gene targets, we compared CXCL13 expression to the most significant transcription factor activity scores in FDC, which express the highest levels of CXCL13 among all cell types in our dataset (Figure 3G-H). Interestingly, the predicted activity of RELB, a downstream mediator of non-canonical NF-κB signaling, had the strongest relationship to CXCL13 expression in FDC. Together, these data suggest that elevated chemokine expression by stromal cells, particularly FDC, may be associated with inflammatory signaling pathways, including non-canonical NF-κB signaling in iMCD-TAFRO.

As CXCL13 is the most dramatically up-regulated chemokine detected in both the blood and lymph node in iMCD-TAFRO, and in other contexts it has been shown to indicate GC activity and CXCR5 up-regulation (18, 19), we profiled CXCR5^+^ cells in circulation and in the lymph node. To determine if high CXCL13 levels were associated with high CXCR5 expression in iMCD-TAFRO, we profiled peripheral blood mononuclear cells (PBMC) from iMCD-TAFRO patients (n=14) and discovered that both T and B lymphocytes expressed, not more, but less CXCR5 compared to healthy donors (n=17). In B cells, we discovered that CXCR5, expressed by nearly all B cells in healthy donors, was downregulated in iMCD-TAFRO during active disease or flare (Figure 4A and B). The percentage of CXCR5^+^ B cells dropped to ∼60% of all B cells and was restored after the flare resolved. The median fluorescence intensity (MFI) of CXCR5 followed the same pattern and was, on average, 3-fold lower in total B cells from iMCD-TAFRO patients during flare, and then rebounded to normal levels in remission (Figure 4C-D). We also profiled T cells in the same patients’ PBMC samples and found that the frequency of cell surface CXCR5-expressing non-naïve CD4^+^ T cells was significantly reduced among circulating T cells during iMCD-TAFRO flare (∼3.0%) compared to healthy donors (∼10%) and restored in remission (∼7%) (Figure 4E-F). The MFI of CXCR5 followed the same pattern and was reduced in non-naïve CD4^+^ T cells from iMCD-TAFRO patients during flare and recovered in remission (Figure 4G-H). Together, these data showing increased levels of chemoattractants and a decrease in cognate receptor expression in circulation could indicate that lymphocytes in the blood expressing these receptors are trafficking out of circulation and into the lymph node or to other regions of heightened chemokine expression.

**Figure 4.**
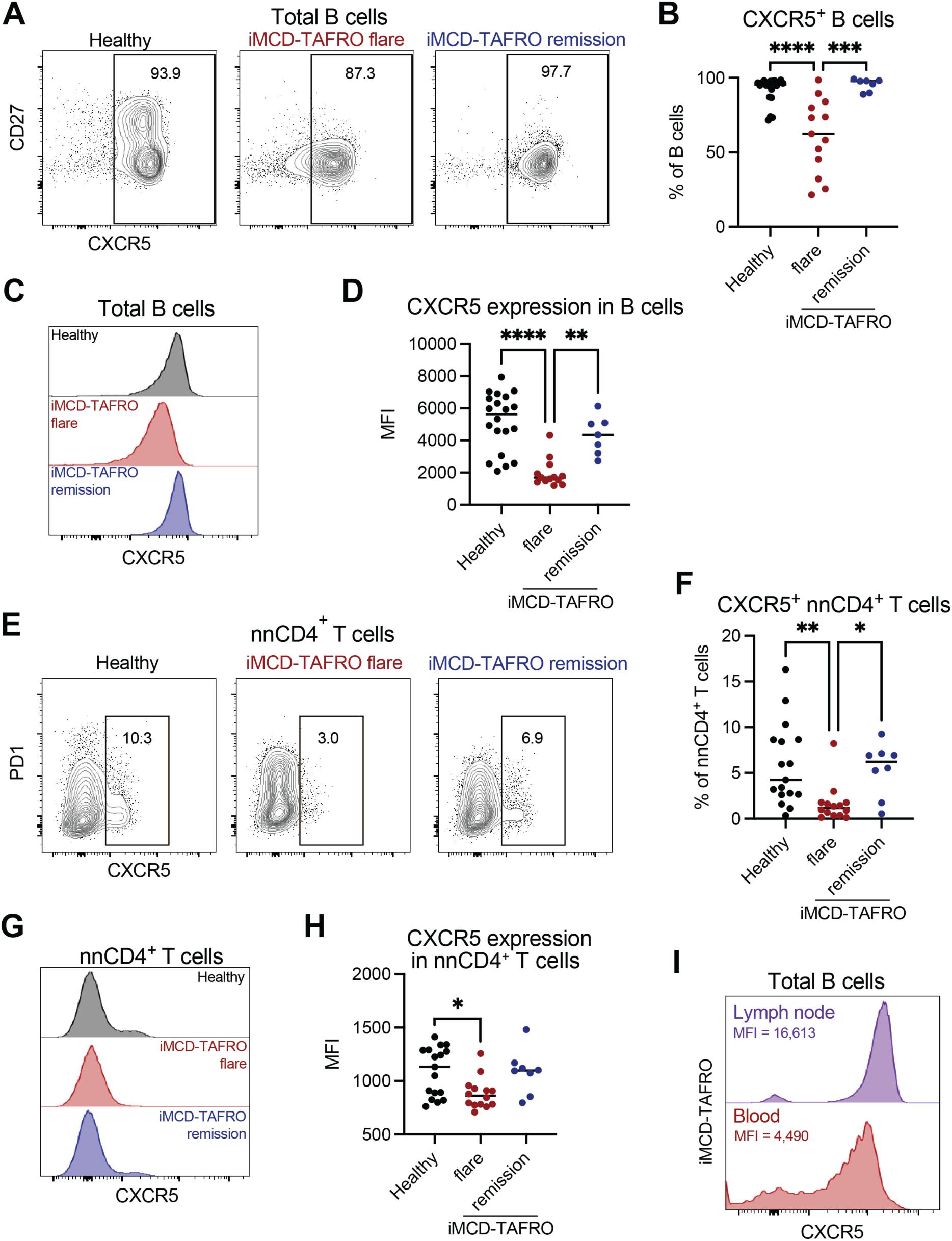
Dysregulated chemokine receptor expression in iMCD-TAFRO. (A) Representative flow cytograms of CXCR5 expression in circulating B cells (CD19^+^) and (B) quantification of the frequency of CXCR5+ cells within the total B cell population within groups. Healthy (n=17); iMCD-TAFRO flare (n=13); iMCD-TAFRO remission (n=7). (C) Representative histogram of CXCR5 expression in B cells in each group. (D) Quantification and comparisons of CXCR5 median fluorescence intensity (MFI) between groups. (E) B cells from the blood and lymph node patient were analyzed for CXCR5 expression (n=1). (F) Representative flow cytogram of non-naïve CD4^+^ T cells (nnCD4^+^) showing the frequency of CXCR5^+^ cells in each group. (G) Comparison of CXCR5^+^ cells within the nnCD4^+^ T cell population per group. Healthy (n=17); iMCD-TAFRO flare (n=14); iMCD-TAFRO remission (n=8). (H) Histogram of CXCR5 expression in nnCD4^+^ T cells and (I) Comparison of CXCR5 MFI between groups. Dots in graphs represent individual patient samples. **p<0.05. **p<0.01. ***p<0.001. ****p<0.001.

To determine if cells with the highest levels of CXCR5 were trafficking out of circulation and into the tissue, we compared CXCR5 expression levels between blood and lymph node. However, this is challenging because CXCR5 profiling at the protein level typically requires fresh tissue, and due to the difficulty in diagnosing iMCD, obtaining fresh lymph node tissue from iMCD-TAFRO patients is rare. Nonetheless, we were able to contemporaneously collect fresh lymph node tissue and peripheral blood from one iMCD-TAFRO patient. Flow cytometry analysis revealed differences in the CXCR5 MFI in lymphocytes. The MFI of CXCR5 in lymph node B cells was 3.7 times higher compared to the same population in the blood (Figure 4I). T cells exhibited a similar pattern of expression, with higher levels in the lymph node compared to the blood, which was expected, as T cells expressing the highest levels of CXCR5^+^ reside in the tissue (Supplemental Figure 3). Although these data are limited to a single patient, they may suggest that immune cells in iMCD-TAFRO that respond to chemoattractants, like CXCL13, mobilize out of circulation and traffic to lymph nodes in iMCD-TAFRO, potentially explaining the lymphadenopathy observed clinically.

Although CXCR5 expression was lower in iMCD-TAFRO lymphocytes in the blood and higher in the lymph node of one iMCD-TAFRO patient, suggesting that T and B lymphocytes traffic to lymph nodes and potentially other SLO, it was still unclear whether the highest expressors of CXCR5 were localizing to the highest regions of CXCL13, the GC, within the tissue in iMCD-TAFRO. We hypothesized that in iMCD-TAFRO, there would be increased numbers of GC B cells and GC-Tfh cells, which express CXCR5 and traffic within the tissue to the follicle via the CXCL13-CXCR5 axis. Using our single-cell RNA seq dataset from lymph nodes, we examined cell populations within the B cell sub-cluster and discovered a lower frequency of cells belonging to what we defined as GC B cells, which expressed *BCL6*, *CD10*, and *AICDA* in iMCD-TAFRO (Figure 5A-B and Supplemental Figure 4A-B). In addition, IF staining using a pan-B cell marker, CD20, also revealed fewer CD20^+^ cells per 100 micron^2^ within GCs in iMCD-TAFRO versus controls (Figure 5C-D).

**Figure 5.**
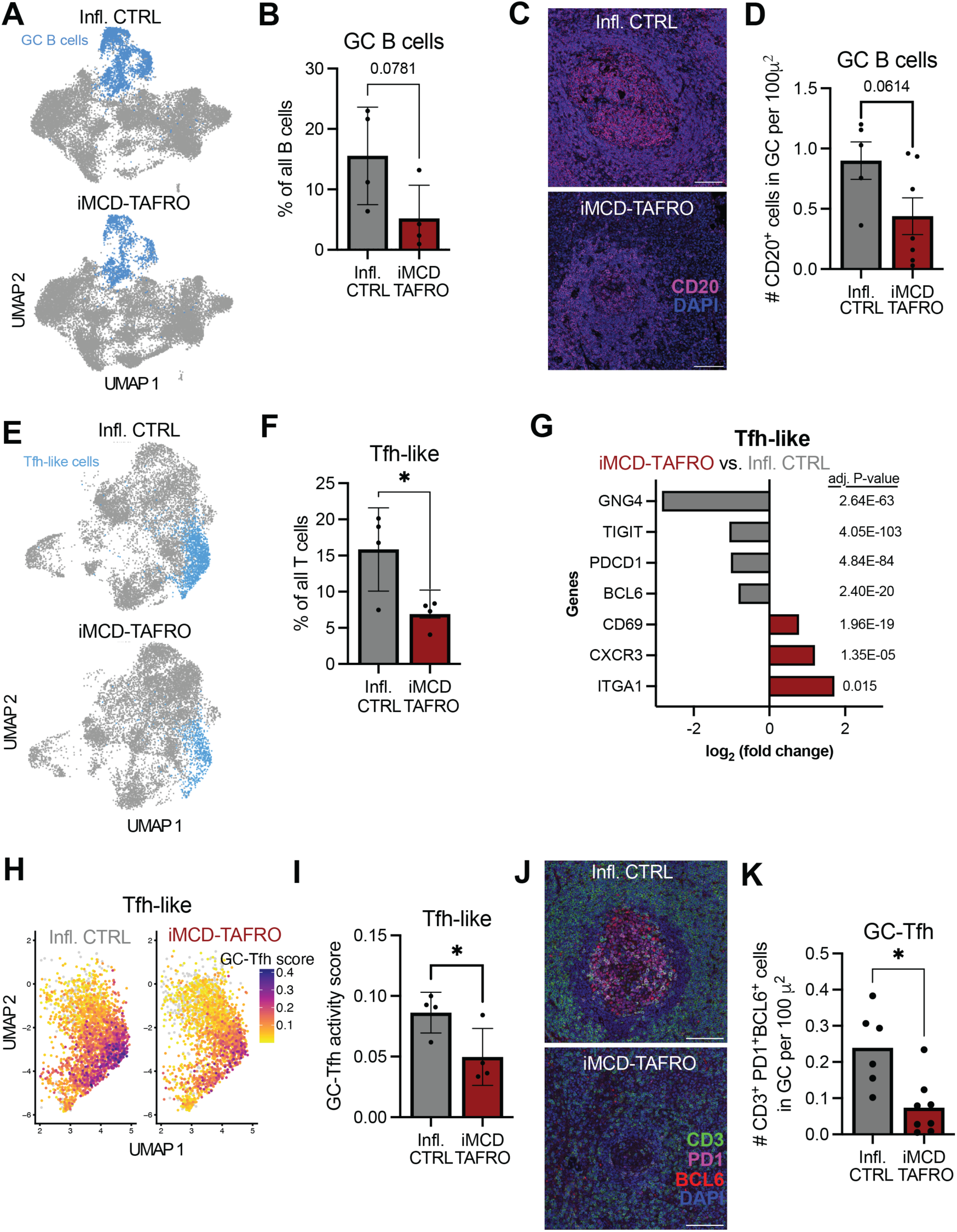
Decreased GC lymphocytes in iMCD-TAFRO lymph nodes. (A) UMAP of B cell sub-cluster highlighting GC B cells. (B) Quantification and comparison of total GC B cells in Infl. CTRL (n=4) and iMCD-TAFRO (n=4). (C) IF images of GCs using B cell marker CD20 and (D) quantification of cells within the GC. p-value is as indicated. (E) UMAP projection of T/NK cell sub-cluster indicating Tfh-like cells in Infl CTRL and iMCD-TAFRO and (F) quantification. (G) Within the Tfh-like cell cluster, differential gene expression comparing iMCD-TAFRO to Infl. CTRL indicates a down-regulation of GC-Tfh genes (grey) and up-regulation of anti-migration-associated genes (red) in iMCD-TAFRO Tfh-like cells. (H) GC-Tfh enrichment scores were applied to the Tfh-like subcluster in both groups. (I) Quantification of the average GC-Tfh activity scores per patient. (J) Representative IF images of GC-Tfh cells that express CD3, PD1, and BCL6 within GCs and (K) quantification. Dots in bar graphs represent individual patient samples. *p<0.05.

We performed a similar analysis for T cells. Using the T cell sub-cluster from our single-cell RNA seq dataset, we defined a Tfh-like cluster based on expression of known Tfh makers including *CXCR5*, *PDCD1*, *BCL6*, and *TOX2* (Supplemental Figure 4C-D). Tfh-like cells were significantly reduced in iMCD-TAFRO lymph nodes relative to inflammatory controls (Figure 5E-F). Further examination of the Tfh-like cluster revealed that genes associated with activation and GC localization, including *PDCD1*, *TIGIT*, and *BCL6* were down-regulated in iMCD-TAFRO (Figure 5G). Furthermore, *GNG4*, recently reported to define activated, GC-localized Tfh in human tonsil ((20), Barnett Dubensky et al., bioRxiv 2025), was among the most significantly downregulated genes in iMCD TAFRO Tfh-like cells (Figure 5G). Using a broader list of features found to be highly up-regulated in GC-localized Tfh (20), we calculated a GC-Tfh enrichment score for each cell in the Tfh-like subcluster (Figure 5H). Tfh-like cells in iMCD-TAFRO subjects exhibited significantly lower GC-Tfh enrichment score, suggesting a reduction in GC localization and functional maturation (Figure 5I). Consistent with our previous findings revealing reduced GC-Tfh in iMCD-TAFRO lymph nodes, expression of GC Tfh-associated genes such as *GNG4* and *TOX2* were significantly down-regulated in the GC region of iMCD-TAFRO lymph nodes in our GeoMx dataset (Figure 2B). To further validate these data, we analyzed IHC slides and performed immunofluorescence (ICC/IF) on lymph node tissue from iMCD-TAFRO patients and inflammatory controls (Figure 5J-K and Supplemental Figure 4E-F). We quantified cells specifically within the GC and discovered that the frequency of total T cells by CD3 staining and number of Tfh (CD3^+^PD1^+^BCL6^+^) per unit area was reduced by approximately 2-fold on average in iMCD-TAFRO lymph nodes. Together, these data suggest that although chemokines including CXCL13 are exceedingly high in the GC of iMCD-TAFRO lymph nodes, CXCR5^+^ subsets such as Tfh do not localize to the GC niche as expected.

Further analysis of the T cell sub-cluster revealed a population of CD4^+^ cells that was approximately 7-times more frequent in iMCD-TAFRO compared to controls (Figure 6A-B). This CD4^+^ T cell population expressed type I interferon (IFN)-responsive genes including *ISG15* and *IFIT1* (Supplemental Figure 4D). In addition, the most up-regulated genes in iMCD-TAFRO Tfh-like cells were IFN-response genes including *IF44*, *IFI44L*, *XAF1* and *MX1*, which are typically associated with type I IFN (Figure 6C) (21, 22). Spatially, the type I IFN gene signature was present in both the GC and IF space (Figure 2B-C). Compared to other CD4^+^ T cells including Tfh-like cells from inflammatory controls, iMCD-TAFRO Tfh-like cells expressed IFNƔ (Figure 6D), a Th1 signature gene, which was especially interesting as type I IFN can restrict Tfh fate and promote Th1 differentiation (23). Furthermore, iMCD-TAFRO Tfh-like cells expressed significantly higher levels of *CD69* and the gene encoding CD49a, *ITGA1*, genes associated with tissue residence (Figure 6D and Figure 5G) (24). Genes associated with migration like *KLF2*, *S1PR1* and *SELL*, which encodes CD62L, were downregulated relative to other CD4^+^ T cell subsets (25, 26). Together, these data suggest that a strong type I IFN signal in iMCD-TAFRO lymph nodes may restrict Tfh differentiation and the ability of T cells to migrate to high CXCL13-expressing GCs in iMCD-TAFRO.

**Figure 6.**
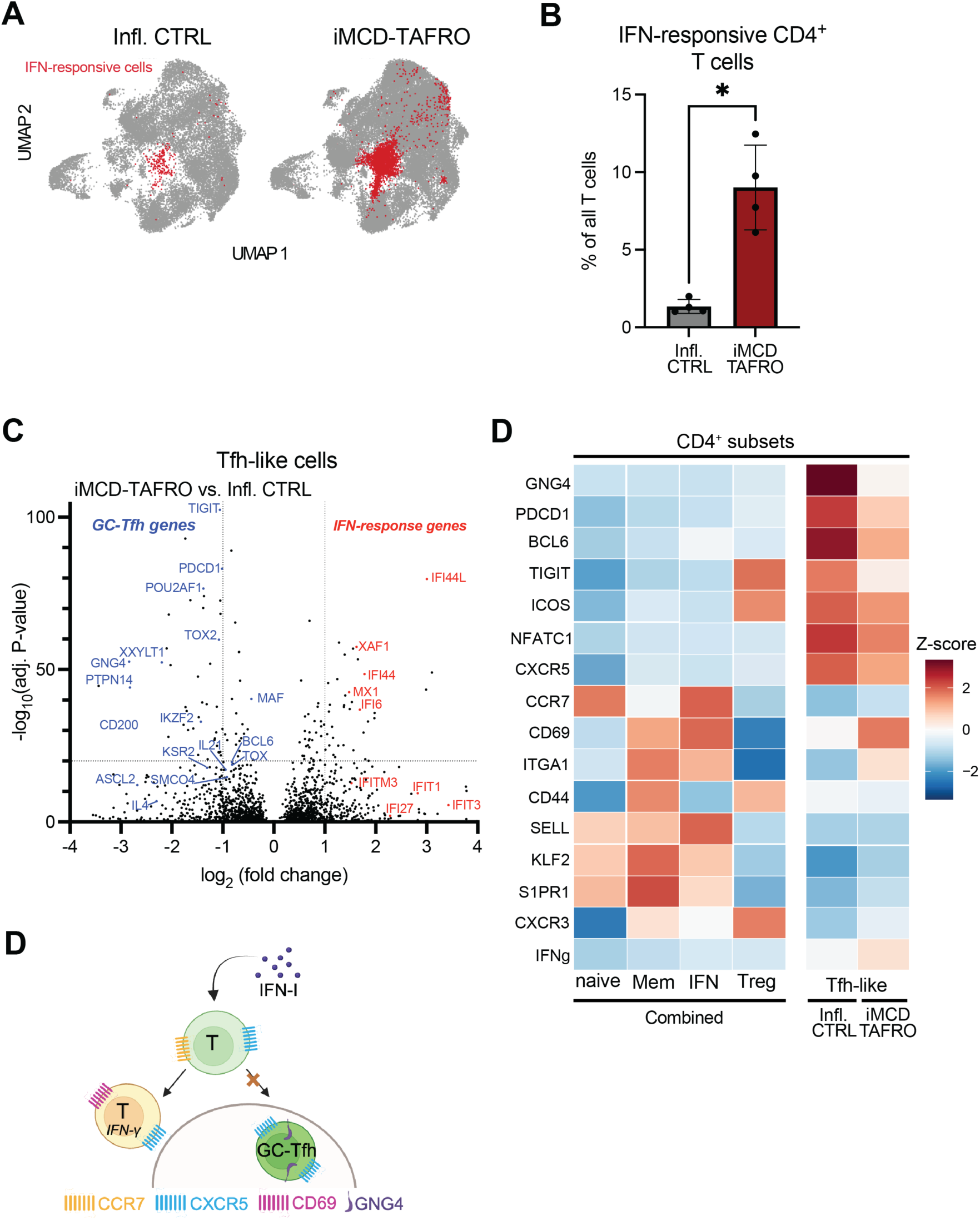
Increased type I IFN gene signatures in CD4^+^ T cells in iMCD-TAFRO lymph nodes. (A) UMAP of T cell sub-cluster highlighting IFN-responsive CD4^+^ T cells in each group. (B) Comparison of the frequency of the IFN-responsive CD4^+^ T cells within the T cell population in iMCD-TAFRO (n=4) and Infl. CTRL patients (n=4). Dots in bar graphs represent individual patient samples. (C) Volcano plot representing differential gene expression between iMCD-TAFRO and Infl. CTRL Tfh-like cells. (D) Heatmap of gene expression within different CD4^+^ T cell populations. Left part of heatmap represents gene expression from all cells in the dataset in a given T cell population and the right side separates gene expression of Tfh-like cells between the two groups. *p<0.05. (E) Proposed model in iMCD-TAFRO lymph nodes in which a strong type I IFN gene signature restricts Tfh production and localization withing the GC. Image generated with Biorender.

Herein, we present data that provides mechanistic insights that may explain the signature lymphadenopathy phenotype of iMCD-TAFRO. We provide evidence that lymph node stromal cells contribute to elevated serum chemokines, including CXCL13, yet are incapable of efficiently attracting key CXCR5^+^ cell types into lymph node GC, a critical step in generating highly specific antibody responses. Our data suggest that T cell differentiation into Tfh is restricted in iMCD-TAFRO, potentially by type I IFN signaling, and retains a residency signature that may prevent efficient GC trafficking. This study indicates two mechanisms that are dysregulated in iMCD-TAFRO—chemotaxis and Tfh differentiation, which warrant further investigation as potential therapeutic targets.

## DISCUSSION

iMCD-TAFRO is a severe and life-threatening disease with limited effective treatments. To reveal novel therapeutic targets, it is critical to understand the pathogenic mechanisms that drive iMCD-TAFRO-specific systemic inflammation, histopathology, and lymphadenopathy. In this study, we discovered that chemokine expression in iMCD-TAFRO associates and potentially drives the generalized lymphadenopathy that is required for diagnosis. We provide evidence that increased lymphocyte trafficking out of circulation and into the lymph node contributes to the enlarged lymph nodes in iMCD-TAFRO. We discovered that heightened chemokine expression from stromal cells in the lymph node from iMCD-TAFRO patients may be responsible for attracting and retaining lymphocytes out of circulation. However, within the tissue, T cells and B cells fail to properly localize to the robust FDC network inside the follicle, thereby failing to form true GC and potentially disrupting and/or preventing normal GC responses.

In health, the GC response is highly coordinated, selecting for B cell clones that bind antigen most successfully to then compete for Tfh help. This tightly regulated process typically yields high-affinity antibodies reactive against non-self antigens. We hypothesize that dysregulated GC formation in iMCD-TAFRO leads to aberrant T:B interactions and promotes autoantibody production, which we previously described in a large proportion of iMCD patients (9). While it is not clear if autoantibody production promotes disease or is a byproduct of dysregulated inflammatory processes in iMCD-TAFRO, it will be interesting to investigate how antigenic stimulation affects GC formation, if at all, in patients with iMCD-TAFRO. Interestingly, despite the dysregulation in GC, iMCD patients can produce antigen-specific antibody responses after vaccination. It remains unclear whether this vaccine-specific antibody production is GC-driven, extrafollicular, or T-cell dependent in this context (27).

As a potential mechanism for GC dysfunction in iMCD-TAFRO, the concurrent loss of GC-Tfh and an increase in IFN-responsive CD4^+^ T cells implicate IFN-driven Tfh cell fate restriction, indicating an immune pathway that may be important for driving non-specific antibody responses. As type I IFN has been shown to impede Tfh differentiation in both mouse and humans (23, 28), our data provide a disease context in which dysregulation of the inflammatory signals provided during T cell priming and differentiation could be pathogenic. Based on their gene expression pattern—namely high *CD69* and high *ITGA1*— the iMCD-TAFRO Tfh-like cells appeared to be less equipped to localize to the follicle and GC. This localization failure is despite the high CXCL13 signal in iMCD-TAFRO, which can also be induced by type I IFN (17). Together, these data suggest that altered T cell differentiation and altered microanatomic organization both add to the immune dysregulation and potential for autoantibody production in iMCD-TAFRO (Supplemental Figure 5).

Our finding that both Tfh gene expression programs, and especially *GNG4* expression, are significantly decreased within dysplastic GC of iMCD-TAFRO patients complements a recent report that *GNG4* marks GC-positioned Tfh (20), which we observe are sparse in iMCD-TAFRO lymph nodes. Our spatial transcriptomic data and single-cell RNA sequencing data show that *GNG4* in iMCD-TAFRO was among the most down-regulated genes in Tfh cells and the GC niche. Furthermore, other known Tfh transcripts that are known to be enriched in GC-phenotype Tfh—including *PDCD1*, *TIGIT*, *TOX*—were also notably down-regulated in iMCD-TAFRO lymph nodes. Indeed, we discovered fewer GC-localized CD3^+^ T cells that expressed PD1 and BCL6 protein in iMCD-TAFRO lymph node tissue compared to controls. While re-analysis of published genome wide association studies data suggest that over-expression of *GNG4* may predispose to autoimmune disorders like rheumatoid and psoriatic arthritis (20, 29, 30), our data provide a distinct hyperinflammatory disease context in which *GNG4* is decreased. Our hypothesis model considers the loss of *GNG4^+^*Tfh to be a consequence of the immune dysregulation in iMCD-TAFRO rather a pathologic driver of the GC dysfunction. Still, the absence of *GNG4* expression despite the presence of GC-phenotype B cells in fine needle aspirates may be a useful measure of lymph node dysfunction and predictive of autoantibody formation.

In addition to proposing a mechanism for non-specific autoantibody production in iMCD-TAFRO, this study also challenges the notion that local expansion of lymphocytes is the primary driver of lymphadenopathy in iMCD. We found that clinical measures of proliferation in the lymph node, as indicated by FDG avidity, are typically unremarkable, with SUVmax values less than 10, and are not positively associated with lymph node size. Instead, we found a positive association between lymph node size and chemokine levels, suggesting that lymphocyte trafficking to the lymph node through chemotaxis contributes to lymph node enlargement in iMCD-TAFRO, potentially promoting disease. Future work investigating the role of chemotaxis in iMCD-TAFRO is needed, as it aims to determine whether it drives disease and thus may serve as a pathogenic mediator and potential therapeutic target.

While our work presented here relies on expression patterns of factors well-known to facilitate cellular trafficking, a model system is needed to evaluate whether chemotaxis is required for disease pathogenesis. Our group recently developed a mouse model that recapitulates several iMCD-TAFRO features, including increased levels of CXCL13 and IL-6, as well as dysregulated GC formation (31). Using this mouse model or another system will be critical to understand the specific factors involved in iMCD-TAFRO and whether their inhibition prevents or mitigates the cytokine storm that can be deadly for patients. This could include testing blocking antibodies against CXCL13 or CXCR5, or preventing their production entirely with a non-canonical NF-κB inhibitor, such as bortezomib. This mouse model could also be used to directly assess CXCR5 protein expression in lymph nodes, which was challenging in this study due to difficulties obtaining fresh tissue from patients and the lack of a suitable CXCR5 antibody that can detect expression within FFPE tissue. Other limitations of this study include our small sample size of iMCD-TAFRO patients, particularly for the transcriptomic analyses. We do not know whether our findings reflect general iMCD-TAFRO pathology or are applicable only to a small subset of patients. Along these lines, it would also be interesting to investigate if these features are specific to iMCD-TAFRO or are consistent among all iMCD subtypes, including iMCD-NOS and iMCD-IPL.

In summary, the work described in this research article provides alternate mechanistic explanations for key pathophysiologic features of iMCD-TAFRO, a severe and life-threatening immunologic disorder with few effective treatment options. Our study suggests that, in iMCD-TAFRO, enhanced lymphocyte trafficking to lymph nodes is associated with lymphadenopathy, and within the tissue, the spatial positioning of lymphocytes is disrupted, potentially contributing to non-specific antibody responses. Future investigation is needed to determine whether these mechanisms are required for iMCD-TAFRO pathogenesis and, therefore, represent prospective therapeutic targets.

## MATERIALS AND METHODS

### Patients

Informed consent was obtained for all patient samples included in this study according to a protocol approved by the University of Pennsylvania Institutional Review Board. Clinical and laboratory data were obtained for iMCD-TAFRO patients enrolled in the ACCELERATE natural history registry (NCT02817997) (32). All iMCD-TAFRO patients were diagnosed based on the iMCD diagnostic criteria (1).

### Serum proteomic analyses

SomaLogic SOMAscan (v4.1) was used to measure 6,408 unique serum analytes from 24 iMCD-TAFRO patients during active disease, defined by 2 or more abnormal clinical laboratory features associated with iMCD used as part of the diagnostic criteria (1), and 15 healthy donors. To identify differentially expressed proteins, raw values for each analyte were log2 transformed and linear regression was applied correcting for age and sex. The Benjamini-Hochberg correction was used to account for multiple comparisons. To determine differentially expressed proteins among cytokines and chemokines, we filtered the analyte panel to 398 unique proteins using two KEGG pathways from msigDB (KEGG_CHEMOKINE_SIGNALING_PATHWAY, KEGG_CYTOKINE_CYTOKINE_RECEPTOR_INTERACTION) (Supplemental Table 1).

### Lymph node size measurements/FDG measurement and quantification

Lymph node size and fluorodeoxyglucose (FDG) measurements were obtained from CT and PET/CT scan records. Short-axis measurements in centimeters were used to determine the size of lymph nodes. Values included the most recent scan obtained within 90 days of the corresponding lymph node excision. For FDG avidity assessment, CT and PET/CT scans were reviewed to identify enlarged lymph nodes and their corresponding FDG avidity, measured by SUV uptake. For each patient, scan dates with lymph nodes greater than 1 cm in size and a measurable SUV value were recorded, with the largest lymph node size noted for each scan date.

### Immunohistochemistry

Lymph node FFPE tissue from iMCD patients were immunohistochemically stained for Ki-67 protein using DAB and hematoxylin and imaged at 20X. A representative cohort of germinal centers, mantle zones, and interfollicular spaces were manually annotated and quantified for each patient. Cells were classified as either positive or negative for Ki-67 using a binary scoring system. For T cell analysis, whole-slide images (.SVS) of CD3-stained FFPE lymph node tissue were imported into QuPath (v0.6.0) for analysis. The stain vector deconvolution tool was used to disable the DAB channel, allowing visualization of the hematoxylin stain only. Lymph node germinal centers (GC), mantle zones (MZ), and the transitional region where the borders of the GC and MZ intersect were manually annotated. Approximately 10 representative annotations were created for each region type and averaged. The positive cell detection tool was then optimized for each image to detect CD3-positive cells. The percentage of CD3-positive cells was calculated for each region type (GC, MZ, and border), and the mean CD3 positivity was calculated for each region.

### Immunofluorescence microscopy and analysis

5-micron sections of FFPE lymph node tissue from iMCD patients and controls were stained in the following manner, following established protocols. Slides were deparaffinized using xylene and ethanol incubations prior to antigen retrieval using antigen retrieval buffer (Abcam; catalog# 93684) under pressure for 3 minutes at 120°C. The tissue was blocked with 0.1% PBS-Tween and 10% FBS. The following primary antibodies were used at the indicated concentrations: anti-CD20 (2ug/mL, Novus catalog#: NBP2-44745), anti-CD3 (10ug/mL, Biorad #MCA1477T), anti-PD-1 (5ug/mL, Thermofisher #14-2798-82), anti-BCL6 (5ug/mL, Cell Signaling #14895T). All secondary antibodies were fluorescently tagged and raised in donkey (1:1000, Thermofisher catalog numbers A-21202, A-78945, and A-31573). Images were obtained using a Zeiss LSM 980 confocal microscope at the UPenn CDB Microscopy Core (RRID SCR_022373) at 40X magnification.

The images were analyzed in the following manner. Nuclei were segmented from the DAPI stain using Cellpose (33). To approximate cytoplasmic boundaries, non-nuclear pixels were assigned to the nearest nucleus. The mean intensity in the nuclear area and peri-nuclear ring were calculated for CD3 and CD20 stains. Cells were determined to be positive or negative for a particular stain by implementing a two-component Gaussian mixture model, where “positive” was defined as the higher-mean component. We then computed the posterior probability curves for both components and used the intersection of these two curves as a threshold cutoff to binarize all cell and cytoplasmic regions into positive and negative categories. This method allowed for adaptation to image-specific artifacts, such as global changes in intensity, while exploiting the knowledge that both positive and negative components for each stain would be present in each image. Germinal centers and mantle zones were identified by manual segmentation. Cells positive for a particular marker were defined by a signal present in the peri-nuclear region for the CD3-specific stain or the CD20-specific stain. For identification of Tfh cells, each signaling channel (CD3, PD-1, BCL6) was processed as described above. The colocalization of these stains was then assessed. Statistical significance was calculated using unpaired T tests with Welsh’s correction when necessary.

### Bruker/Nanostring GeoMx

Histology sections (5 um) of FFPE tissue from each sample was cut fresh and stained with antibodies against CD3 (Novus # NBP2-54392AF532) and CD20 (Novus # NBP2-47840AF594) and nuclear dye Sytox13. Slides were prepped according to manufacturer’s instructions (Bruker/Nanostring GeoMx) as described previously (34). Normalized and pooled GeoMx samples were quantified and sequenced on the Illumina NovaSeq 6000 platform on an S1-100 flow cell. Paired-end sequencing was performed as recommended: 27 × 8 × 8 × 27. The final pool was denatured and diluted to a loading concentration of 250pM. PhiX control was added at a concentration of 1%.

Digital count conversion (DCC) files were downloaded from the Bruker/Nanostring Digital Spatial Profiler (DSP). DCC files were read into the R compute environment using the GeoMxTools package (35). Raw counts were log transformed and background corrected using the default SpatialDecon preprocessing workflow to stabilize variance across segments. Several quality control parameters were used to select high quality segments including, minimum number of reads > 1000, minimum percent of reads trimmed > 80%, minimum percent of reads stitched > 80%, minimum percent of reads aligned >75%, minimum sequencing saturation > 50%, minimum negative control counts >10, minimum number of nuclei estimated > 100, and minimum segment area > 1000. Flagged segments were then removed from further analysis. After filtering, 176 segments were used for spatial analysis. We used upper-quartile (Q3) normalization to compute per-segment scaling factors ensuring that differences in library size did not obscure biological variation in downstream visualization and differential expression analysis. A linear mixed model was used to identify differentially expressed genes between groups of segments. To account for multiple comparisons, the false discovery rate (FDR) method was used.

To perform spatial deconvolution of each segment, the SpatialDecon package was used (36). Cell type proportion scores were scaled across segments to allow comparison between groups and to reduce technical variation. We used the annotated 10X Genomics’ FLEX dataset described below as a reference to deconvolute each segment into estimated cell type proportions.

### Single-cell RNA sequencing

Patient samples were processed according to 10X Genomics’ standard FLEX protocol. Briefly, FFPE tissues were cut into 25um sections and dissociated using a gentleMACS Dissociator (Miltenyi Biotech) followed by the GEM-X-Flex v2 protocol (10X Genomics). Eight samples were partitioned into two pools (Pool A, 4 samples and Pool B, 4 samples). The final pools were quantified and sequenced on the Illumina NovaSeq 6000 platform using an S1-100 flow cell. Paired-end sequencing was performed according to 10X Genomics’ recommendations: 28 × 10 × 10 × 90. The final pool was denatured and diluted to a loading concentration of 230pM. PhiX control was added at a concentration of 1%.

The Chromium Human Transcriptome Probe Set (v1.0.1) was used to map sequencing reads to the GRCh38 human reference using the CellRanger (v7.2) pipeline. Filtered feature barcode matrices were read into the R compute environment (v. 4.5.2) using the Seurat library (37). Multiple filtering criteria were applied to each sample, including min.cells = 3, 200 < nFeature < 5000, and percent mitochondrial gene expression < 10%. After gene and cell quality level filtering, the final dataset consisted of 18,054 genes expressed in 168,504 cells. Gene expression data was normalized and scaled, and integration was performed using the harmony package (38). Cell cycle score, mitochondrial gene expression, and Interferon alpha and gamma signature scores were regressed out of the principal component analysis (PCA) before downstream clustering. Initial clustering resulted in 53 cell clusters. Differentially expressed gene markers per cluster were produced using the FindAllMarkers function and using the MAST test (39). Top differentially expressed genes were used for initial annotation. Based on increased expression of cell type marker genes, cells were subsetted into 4 subpopulations -- non-hematopoietic/stromal cells, T and NK cells, B cells, and myeloid cells. These subpopulations were re-clustered and manually annotated using canonical cell type markers. For single cell gene set enrichment analysis, the escape package was used with the hallmark gene sets from the msigdb repository (40). To determine the GC-Tfh signature score, the AddModuleScore function in Seurat was used to calculate a composite score of 232 genes from (20). To perform transcription factor (TF) analysis, the DoRothEA package was used along with the statistical package VIPER for evaluation (41, 42). To account for multiple comparisons in all statistical tests, a Bonferroni correction was applied.

### Flow cytometry

PBMCs were thawed, washed, and rested for 2h in R10 media (RPMI 1640 supplemented with 2 mM L-glutamine, 100 U/ml of penicillin and streptomycin, and 10% FBS -Gemini-). Whole blood and lymph node samples collected on the same day were processed as described previously (43). After washing, cells were pre-stained for chemokine receptors for 15 min at 37°C 5% CO2. All following incubations were performed at room temperature. Cells were stained with a viability dye (Live/Dead Fixable Aqua) for 10 minutes, followed by a 20-minute staining of surface markers. Antibodies were diluted in equal parts of FACS buffer (0.1% sodium azide and 1% bovine serum albumin in 1X PBS) and BD Brilliant Stain buffer (BD Biosciences). Then, cells were washed and fixed/permeabilized using the FoxP3 Transcription Factor Buffer Kit (eBioscience), following the manufacturer’s instructions. Intracellular staining was performed by adding the antibody cocktail prepared in 1X Permwash buffer for 1 hour at 37°C. Stained cells were washed and fixed in PBS containing 1% paraformaldehyde (Sigma-Aldrich, St. Luis, MO), and stored at 4°C in the dark until acquisition. The antibodies used are listed in Supplemental Table 2. All flow cytometry data were collected on a BD FACSymphony A5 cytometer (BD Biosciences), and data were analyzed using FlowJo™ v10.8.1 Software (BD Life Sciences).

### Statistics

Unpaired, student’s t-test were performed when comparisons were made between 2 groups using Prism 10 software (Graphpad). Welch’s correction was applied when the calculated F-test provided significantly different variances among groups. For serum proteomics comparing chemokines between inflammatory diseases, one-way ANOVA was used for multiple comparisons and a Bonferroni correction was applied. For other statistical tests used, see other sections above.

## Acknowledgements

The authors would like to thank the patients and their families for their continued support of this work and participation in our ACCELERATE registry. From the University of Pennsylvania, we thank the Molecular Pathology and Imaging Core (RRID: SCR_022420) where we performed the tissue deparaffinization and antigen retrieval for our IF studies, the Cell and Developmental Microscopy Core (RRID: SCR_022373) for access to their imaging microscopes, the Penn Cytomics and Cell Sorting Shared Resource Laboratory (RRID: SCR_022376) for access to the flow cytometers, and the Human Immunology Core (RRID:SCR_022380) for processing of patient serum and PBMC samples. We also are grateful for the CHOP Single-Cell Core for assisting us with the Bruker/Nanostring GeoMx experiment, the CHOP High-Throughput Sequencing Core Laboratory where we performed all the sequencing in this study, and the CHOP Pathology Core Laboratory (RRID:SCR_009726) for tissue processing and IHC studies. Lastly, we thank Dorottya Laczko and Taku Kambayashi for their helpful discussions regarding Tfh cells, models of iMCD, and broader implications of our study.

## Author Contributions

MDM and DCF developed the study methodology. MDM, KSF, and MBP performed the experiments and MDM, MVG, KSF, MBP, AHI, CLML, JMZ, and IDM analyzed data. MDM, MVG, KSF, MBP, SBD, DAO, LAV, MRB, and DCF interpreted the data. MDM and DCF prepared the initial manuscript. The final manuscript was reviewed, edited, and approved by all the authors.

## Data Availability

Transcriptomics’ datasets including Bruker/Nanostring GeoMx and 10X Genomics FLEX were deposited to Gene Expression Omnibus and available through accession number GSE######.

## References

1. Fajgenbaum DC, Uldrick TS, Bagg A, Frank D, Wu D, Srkalovic G, Simpson D, Liu AY, Menke D, Chandrakasan S, Lechowicz MJ, Wong RS, Pierson S, Paessler M, Rossi JF, Ide M, Ruth J, Croglio M, Suarez A, Krymskaya V, Chadburn A, Colleoni G, Nasta S, Jayanthan R, Nabel CS, Casper C, Dispenzieri A, Fossa A, Kelleher D, Kurzrock R, Voorhees P, Dogan A, Yoshizaki K, van Rhee F, Oksenhendler E, Jaffe ES, Elenitoba-Johnson KS, Lim MS. 2017. International, evidence-based consensus diagnostic criteria for HHV-8-negative/idiopathic multicentric Castleman disease. Blood 129: 1646–57

2. Fajgenbaum DC. 2018. Novel insights and therapeutic approaches in idiopathic multicentric Castleman disease. Blood 132: 2323–30

3. Pierson SK, Lim MS, Srkalovic G, Brandstadter JD, Sarmiento Bustamante M, Shyamsundar S, Mango N, Lavery C, Austin B, Alapat D, Lechowicz MJ, Bagg A, Li H, Casper C, van Rhee F, Fajgenbaum DC. 2023. Treatment consistent with idiopathic multicentric Castleman disease guidelines is associated with improved outcomes. Blood Adv 7: 6652–64

4. Nishikori A, Nishimura MF, Nishimura Y, Otsuka F, Maehama K, Ohsawa K, Momose S, Nakamura N, Sato Y. 2022. Idiopathic Plasmacytic Lymphadenopathy Forms an Independent Subtype of Idiopathic Multicentric Castleman Disease. Int J Mol Sci 23

5. Bustamante MS, Pierson SK, Ren Y, Bagg A, Brandstadter JD, Srkalovic G, Mango N, Alapat D, Lechowicz MJ, Li H, Van Rhee F, Lim MS, Fajgenbaum DC. 2024. Longitudinal, natural history study reveals the disease burden of idiopathic multicentric Castleman disease. Haematologica 109: 2196–206

6. Kawabata H, Takai K, Kojima M, Nakamura N, Aoki S, Nakamura S, Kinoshita T, Masaki Y. 2013. Castleman-Kojima disease (TAFRO syndrome) : a novel systemic inflammatory disease characterized by a constellation of symptoms, namely, thrombocytopenia, ascites (anasarca), microcytic anemia, myelofibrosis, renal dysfunction, and organomegaly : a status report and summary of Fukushima (6 June, 2012) and Nagoya meetings (22 September, 2012). J Clin Exp Hematop 53: 57–61

7. Iwaki N, Sato Y, Takata K, Kondo E, Ohno K, Takeuchi M, Orita Y, Nakao S, Yoshino T. 2013. Atypical hyaline vascular-type castleman’s disease with thrombocytopenia, anasarca, fever, and systemic lymphadenopathy. J Clin Exp Hematop 53: 87–93

8. Miller I, Mumau MD, Shyamsundar S, Sarmiento Bustamante M, Horna P, Gonzalez MV, Fajgenbaum DC. 2025. No evidence for active viral infection in unicentric and idiopathic multicentric Castleman disease by Viral-Track analysis. Sci Rep 15: 1676

9. Feng A, Gonzalez MV, Kalaycioglu M, Yin X, Mumau M, Shyamsundar S, Bustamante MS, Chang SE, Dhingra S, Dodig-Crnkovic T, Schwenk JM, Garg T, Yoshizaki K, van Rhee F, Fajgenbaum DC, Utz PJ. 2025. Common connective tissue disorder and anti-cytokine autoantibodies are enriched in idiopathic multicentric castleman disease patients. Front Immunol 16: 1528465

10. Fajgenbaum DC, Langan RA, Japp AS, Partridge HL, Pierson SK, Singh A, Arenas DJ, Ruth JR, Nabel CS, Stone K, Okumura M, Schwarer A, Jose FF, Hamerschlak N, Wertheim GB, Jordan MB, Cohen AD, Krymskaya V, Rubenstein A, Betts MR, Kambayashi T, van Rhee F, Uldrick TS. 2019. Identifying and targeting pathogenic PI3K/AKT/mTOR signaling in IL-6-blockade-refractory idiopathic multicentric Castleman disease. J Clin Invest 129: 4451–63

11. Mumau MD, Gonzalez MV, Ma C, Irvine AH, Sarmiento Bustamante M, Shyamsundar S, Chen LYC, Koslicki D, Fajgenbaum DC. 2025. Identifying and Targeting TNF Signaling in Idiopathic Multicentric Castleman’s Disease. N Engl J Med 392: 616–8

12. Pierson SK, Katz L, Williams R, Mumau M, Gonzalez M, Guzman S, Rubenstein A, Oromendia AB, Beineke P, Fossa A, van Rhee F, Fajgenbaum DC. 2022. CXCL13 is a predictive biomarker in idiopathic multicentric Castleman disease. Nat Commun 13: 7236

13. Pierson SK, Shenoy S, Oromendia AB, Gorzewski AM, Langan Pai RA, Nabel CS, Ruth JR, Parente SAT, Arenas DJ, Guilfoyle M, Reddy M, Weinblatt M, Shadick N, Bower M, Pria AD, Masaki Y, Katz L, Mezey J, Beineke P, Lee D, Tendler C, Kambayashi T, Fossa A, van Rhee F, Fajgenbaum DC. 2021. Discovery and validation of a novel subgroup and therapeutic target in idiopathic multicentric Castleman disease. Blood Adv 5: 3445–56

14. Pierson SK, Stonestrom AJ, Shilling D, Ruth J, Nabel CS, Singh A, Ren Y, Stone K, Li H, van Rhee F, Fajgenbaum DC. 2018. Plasma proteomics identifies a ’chemokine storm’ in idiopathic multicentric Castleman disease. Am J Hematol 93: 902–12

15. Smith D, Eichinger A, Fennell E, Xu-Monette ZY, Rech A, Wang J, Esteva E, Seyedian A, Yang X, Zhang M, Martinez D, Tan K, Luo M, Young KJ, Murray PG, Park C, Reizis B, Pillai V. 2025. Spatial and single cell mapping of castleman disease reveals key stromal cell types and cytokine pathways. Nat Commun 16: 6009

16. Dejardin E, Droin NM, Delhase M, Haas E, Cao Y, Makris C, Li ZW, Karin M, Ware CF, Green DR. 2002. The lymphotoxin-beta receptor induces different patterns of gene expression via two NF-kappaB pathways. Immunity 17: 525–35

17. Denton AE, Innocentin S, Carr EJ, Bradford BM, Lafouresse F, Mabbott NA, Morbe U, Ludewig B, Groom JR, Good-Jacobson KL, Linterman MA. 2019. Type I interferon induces CXCL13 to support ectopic germinal center formation. J Exp Med 216: 621–37

18. Havenar-Daughton C, Lindqvist M, Heit A, Wu JE, Reiss SM, Kendric K, Belanger S, Kasturi SP, Landais E, Akondy RS, McGuire HM, Bothwell M, Vagefi PA, Scully E, Investigators IPCP, Tomaras GD, Davis MM, Poignard P, Ahmed R, Walker BD, Pulendran B, McElrath MJ, Kaufmann DE, Crotty S. 2016. CXCL13 is a plasma biomarker of germinal center activity. Proc Natl Acad Sci U S A 113: 2702–7

19. Vella LA, Buggert M, Manne S, Herati RS, Sayin I, Kuri-Cervantes L, Bukh Brody I, O’Boyle KC, Kaprielian H, Giles JR, Nguyen S, Muselman A, Antel JP, Bar-Or A, Johnson ME, Canaday DH, Naji A, Ganusov VV, Laufer TM, Wells AD, Dori Y, Itkin MG, Betts MR, Wherry EJ. 2019. T follicular helper cells in human efferent lymph retain lymphoid characteristics. J Clin Invest 129: 3185–200

20. 20. Barnett Dubensky S, Zhu Y, Gallagher M, Kumashie KG, Lu T, Tedesco J, De Luna N, Premo K, Qi Y, Rachimi S, Cabrera EC, Fulmer B, Meremikwu IC, Carter A, Henrickson SE, Romberg N, Baxter AE, Vella LA. 2025. Multimodal analysis defines GNG4 as a distinguishing feature of germinal center-positioned CD4 T follicular helper cells in humans bioRxiv

21. Liu SY, Sanchez DJ, Aliyari R, Lu S, Cheng G. 2012. Systematic identification of type I and type II interferon-induced antiviral factors. Proc Natl Acad Sci U S A 109: 4239–44

22. Moss N, Sakai C, Kaul SN, Graybuck LT, Rachid Zaim S, Angus-Hill ML, He YD, Layton ED, Bouvatte P, Wittig PJ, La France CM, Peng T, Glass MC, Krishnan U, Chander A, Kawelo EK, Garber J, Reading J, Anover-Sombke SD, Kwok M, Green DJ, Goldrath AW, Sigvardsson M, Skene PJ, Li X-j, Torgerson TR, Kuan EL. 2025. Dissecting type I and II interferon impacts on human immune cells in disease by a cell type-specific interferon response atlas. bioRxiv: 2025.12.02.691676

23. Ray JP, Marshall HD, Laidlaw BJ, Staron MM, Kaech SM, Craft J. 2014. Transcription factor STAT3 and type I interferons are corepressive insulators for differentiation of follicular helper and T helper 1 cells. Immunity 40: 367–77

24. Thome JJ, Yudanin N, Ohmura Y, Kubota M, Grinshpun B, Sathaliyawala T, Kato T, Lerner H, Shen Y, Farber DL. 2014. Spatial map of human T cell compartmentalization and maintenance over decades of life. Cell 159: 814–28

25. Kumar BV, Ma W, Miron M, Granot T, Guyer RS, Carpenter DJ, Senda T, Sun X, Ho SH, Lerner H, Friedman AL, Shen Y, Farber DL. 2017. Human Tissue-Resident Memory T Cells Are Defined by Core Transcriptional and Functional Signatures in Lymphoid and Mucosal Sites. Cell Rep 20: 2921–34

26. Steinbach K, Vincenti I, Merkler D. 2018. Resident-Memory T Cells in Tissue-Restricted Immune Responses: For Better or Worse? Front Immunol 9: 2827

27. Shyamsundar S, Pierson SK, Connolly CM, Teles M, Segev DL, Werbel WA, van Rhee F, Casper C, Brandstadter JD, Noy A, Fajgenbaum DC. 2024. Castleman disease patients report mild COVID-19 symptoms and mount a humoral response to SARS-CoV-2 vaccination. Blood Neoplasia 1

28. Tanemura S, Tsujimoto H, Seki N, Kojima S, Miyoshi F, Sugahara K, Yoshimoto K, Suzuki K, Kaneko Y, Chiba K, Takeuchi T. 2022. Role of interferons (IFNs) in the differentiation of T peripheral helper (Tph) cells. Int Immunol 34: 519–32

29. Ishigaki K, Sakaue S, Terao C, Luo Y, Sonehara K, Yamaguchi K, Amariuta T, Too CL, Laufer VA, Scott IC, Viatte S, Takahashi M, Ohmura K, Murasawa A, Hashimoto M, Ito H, Hammoudeh M, Emadi SA, Masri BK, Halabi H, Badsha H, Uthman IW, Wu X, Lin L, Li T, Plant D, Barton A, Orozco G, Verstappen SMM, Bowes J, MacGregor AJ, Honda S, Koido M, Tomizuka K, Kamatani Y, Tanaka H, Tanaka E, Suzuki A, Maeda Y, Yamamoto K, Miyawaki S, Xie G, Zhang J, Amos CI, Keystone E, Wolbink G, van der Horst-Bruinsma I, Cui J, Liao KP, Carroll RJ, Lee HS, Bang SY, Siminovitch KA, de Vries N, Alfredsson L, Rantapaa-Dahlqvist S, Karlson EW, Bae SC, Kimberly RP, Edberg JC, Mariette X, Huizinga T, Dieude P, Schneider M, Kerick M, Denny JC, BioBank Japan P, Matsuda K, Matsuo K, Mimori T, Matsuda F, Fujio K, Tanaka Y, Kumanogoh A, Traylor M, Lewis CM, Eyre S, Xu H, Saxena R, Arayssi T, Kochi Y, Ikari K, Harigai M, Gregersen PK, Yamamoto K, Louis Bridges S, Jr., Padyukov L, Martin J, Klareskog L, Okada Y, Raychaudhuri S. 2022. Multi-ancestry genome-wide association analyses identify novel genetic mechanisms in rheumatoid arthritis. Nat Genet 54: 1640–51

30. Verma A, Huffman JE, Rodriguez A, Conery M, Liu M, Ho YL, Kim Y, Heise DA, Guare L, Panickan VA, Garcon H, Linares F, Costa L, Goethert I, Tipton R, Honerlaw J, Davies L, Whitbourne S, Cohen J, Posner DC, Sangar R, Murray M, Wang X, Dochtermann DR, Devineni P, Shi Y, Nandi TN, Assimes TL, Brunette CA, Carroll RJ, Clifford R, Duvall S, Gelernter J, Hung A, Iyengar SK, Joseph J, Kember R, Kranzler H, Kripke CM, Levey D, Luoh SW, Merritt VC, Overstreet C, Deak JD, Grant SFA, Polimanti R, Roussos P, Shakt G, Sun YV, Tsao N, Venkatesh S, Voloudakis G, Justice A, Begoli E, Ramoni R, Tourassi G, Pyarajan S, Tsao P, O’Donnell CJ, Muralidhar S, Moser J, Casas JP, Bick AG, Zhou W, Cai T, Voight BF, Cho K, Gaziano JM, Madduri RK, Damrauer S, Liao KP. 2024. Diversity and scale: Genetic architecture of 2068 traits in the VA Million Veteran Program. Science 385: eadj1182

31. Laczko D, Tsao PY, Pai RA, Zinski J, Gonzalez MV, Mumau M, Raabe TD, Muramatsu H, Pardi N, Kambayashi T, Fajgenbaum DC. 2025. A patient-derived CABIN1 mutation recapitulates features of idiopathic multicentric Castleman disease in a mouse model. Blood Adv

32. Pierson SK, Khor JS, Ziglar J, Liu A, Floess K, NaPier E, Gorzewski AM, Tamakloe MA, Powers V, Akhter F, Haljasmaa E, Jayanthan R, Rubenstein A, Repasky M, Elenitoba-Johnson K, Ruth J, Jacobs B, Streetly M, Angenendt L, Patier JL, Ferrero S, Zinzani PL, Terriou L, Casper C, Jaffe E, Hoffmann C, Oksenhendler E, Fossa A, Srkalovic G, Chadburn A, Uldrick TS, Lim M, van Rhee F, Fajgenbaum DC. 2020. ACCELERATE: A Patient-Powered Natural History Study Design Enabling Clinical and Therapeutic Discoveries in a Rare Disorder. Cell Rep Med 1: 100158

33. Stringer C, Wang T, Michaelos M, Pachitariu M. 2021. Cellpose: a generalist algorithm for cellular segmentation. Nat Methods 18: 100–6

34. Merritt CR, Ong GT, Church SE, Barker K, Danaher P, Geiss G, Hoang M, Jung J, Liang Y, McKay-Fleisch J, Nguyen K, Norgaard Z, Sorg K, Sprague I, Warren C, Warren S, Webster PJ, Zhou Z, Zollinger DR, Dunaway DL, Mills GB, Beechem JM. 2020. Multiplex digital spatial profiling of proteins and RNA in fixed tissue. Nat Biotechnol 38: 586–99

35. Griswold M, Ortogero N, Yang Z, Vitancol R, Henderson D. 2025. GeomxTools: NanoString GeoMx Tools..

36. Danaher P, Kim Y, Nelson B, Griswold M, Yang Z, Piazza E, Beechem JM. 2022. Advances in mixed cell deconvolution enable quantification of cell types in spatial transcriptomic data. Nat Commun 13: 385

37. Hao Y, Stuart T, Kowalski MH, Choudhary S, Hoffman P, Hartman A, Srivastava A, Molla G, Madad S, Fernandez-Granda C, Satija R. 2024. Dictionary learning for integrative, multimodal and scalable single-cell analysis. Nat Biotechnol 42: 293–304

38. Korsunsky I, Millard N, Fan J, Slowikowski K, Zhang F, Wei K, Baglaenko Y, Brenner M, Loh PR, Raychaudhuri S. 2019. Fast, sensitive and accurate integration of single-cell data with Harmony. Nat Methods 16: 1289–96

39. Finak G, McDavid A, Yajima M, Deng J, Gersuk V, Shalek AK, Slichter CK, Miller HW, McElrath MJ, Prlic M, Linsley PS, Gottardo R. 2015. MAST: a flexible statistical framework for assessing transcriptional changes and characterizing heterogeneity in single-cell RNA sequencing data. Genome Biol 16: 278

40. Borcherding N, Vishwakarma A, Voigt AP, Bellizzi A, Kaplan J, Nepple K, Salem AK, Jenkins RW, Zakharia Y, Zhang W. 2021. Mapping the immune environment in clear cell renal carcinoma by single-cell genomics. Commun Biol 4: 122

41. Holland CH, Tanevski J, Perales-Paton J, Gleixner J, Kumar MP, Mereu E, Joughin BA, Stegle O, Lauffenburger DA, Heyn H, Szalai B, Saez-Rodriguez J. 2020. Robustness and applicability of transcription factor and pathway analysis tools on single-cell RNA-seq data. Genome Biol 21: 36

42. Alvarez MJ, Shen Y, Giorgi FM, Lachmann A, Ding BB, Ye BH, Califano A. 2016. Functional characterization of somatic mutations in cancer using network-based inference of protein activity. Nat Genet 48: 838–47

43. Kuri-Cervantes L, Pampena MB, Betts MR. 2020. Phenotypic Characterization of SLe(x)+ and CLA+ CD4+ T Cells. STAR Protoc 1: 100154

