## Supplemental Figures for "Dysregulated lymphocyte localization in idiopathic multicentric Castleman disease"

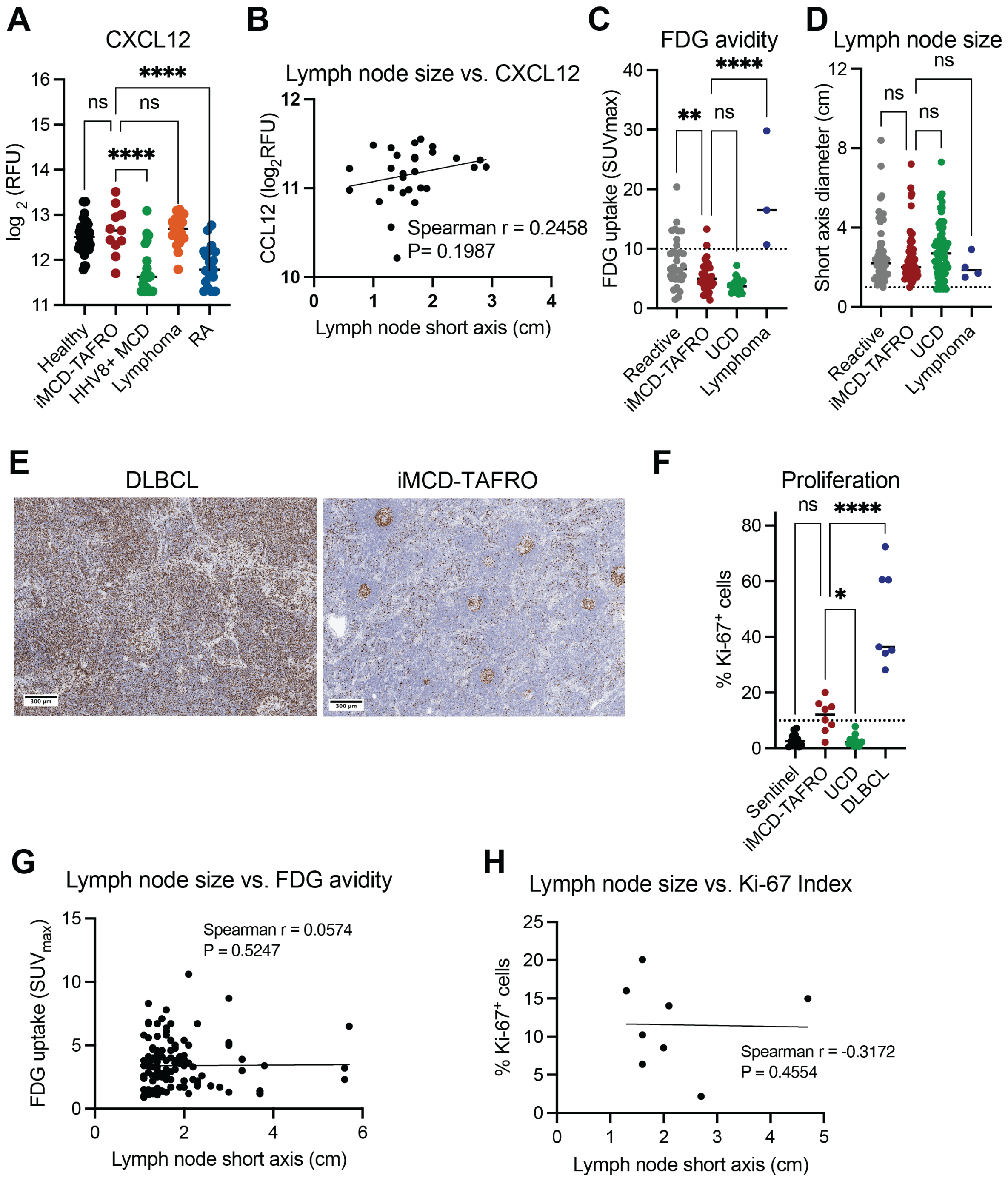


**Supplemental Figure 1. Proliferation is unremarkable in iMCD-TAFRO lymph node tissue.** (A) CXCL12 levels were compared between iMCD-TAFRO and other inflammatory diseases (Healthy, n=42; iMCD-TAFRO, n=11; human herpesvirus-8 associated MCD ($HHV8^+$ MCD), n=20; Hodgkin's lymphoma (Lymphoma), n=20; Rheumatoid arthritis (RA), n=19). (B) Levels of

CXCL12 were plotted against lymph node sizes taken from clinical radiological data. (C) Fluorodeoxyglucose (FDG) uptake from clinical radiology data was compared between iMCD-TAFRO and control lymph nodes (Reactive, n=33; iMCD-TAFRO, n=36, Unicentric Castleman disease (UCD) n=18; lymphoma, n= 3). Dotted line at $SUV_{max}$ of 10 indicates the cut-off for aggressive vs. indolent disease [44]. (D) Short axis lymph node size was compared between iMCD-TAFRO and control lymph nodes (Reactive, n=53; iMCD-TAFRO; n=59, UCD, n=65; lymphoma, n= 3). Dotted line at y=1cm represents lower end of what is considered lymphadenopathy. (E) Representative images of immunohistochemistry against Ki-67 in Diffuse large B-cell lymphoma (DLBCL) and iMCD-TAFRO. (F) Quantification of frequency of Ki-67$^{+}$ cells in iMCD-TAFRO and control lymph nodes (Sentinel, n=11; iMCD-TAFRO, n=8; UCD, n=13; DLBCL n=7). A frequency at or above 10% Ki-67$^{+}$ is considered abnormal indicated by the dotted line. (G) FDG uptake was plotted against the largest lymph node measured on the date closest to the blood collection date used in chemokine assessment. (H) Percentage of Ki-67 positive cells from immunohistochemistry analysis was plotted against lymph node sizes taken from clinical radiology data. *$p<0.05$. **$p<0.01$ ***$p<0.001$ ****$p<0.0001$.

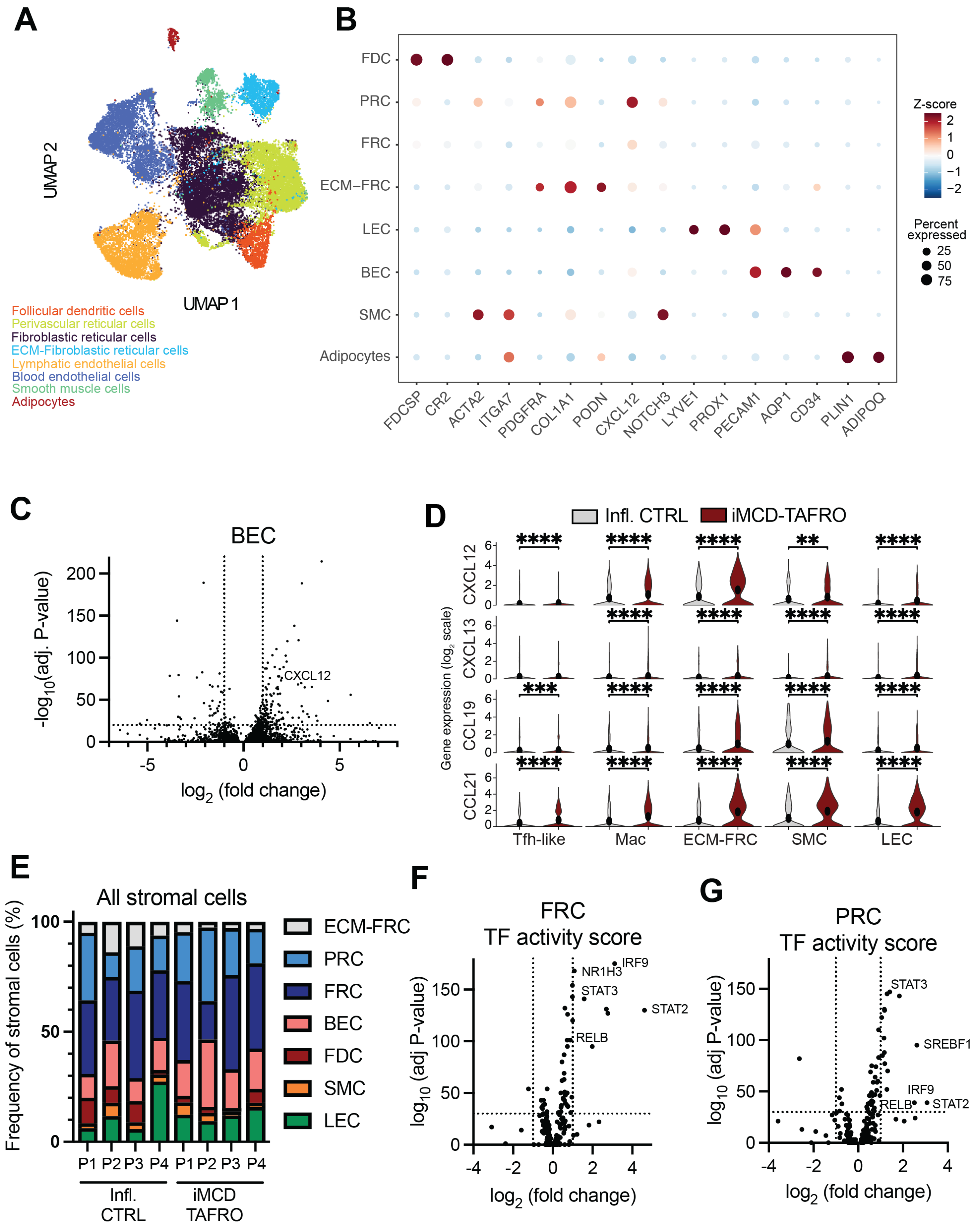


**Supplemental Figure 2. Stromal cell characterization in iMCD lymph nodes.**

(A) Uniform manifold approximation projection (UMAP) of scRNAseq non-hematopoietic/stromal cell sub-cluster. (B) Dot plot showing gene expression of stromal cell lineage markers in each stromal cell sub-cluster. The size of each dot represents the percent of cells expressing the gene and color representing Z score. (C) Volcano plot comparing iMCD-TAFRO vs. Infl. CTRL Blood Endothelial cells (BECs) with CXCL12 highlighted as one of the top differentially-expressed genes. (D) Violin plots comparing CXCL12, CXCL13, CCL19, and CCL21 in hematopoietic (T follicular helper (Tfh) -like (Tfh) and macrophages (Mac), see Supplemental Figure 3) and non-hematopoietic/stromal cell types (extracellular matrix-producing fibroblastic reticular cells (ECM-FRC), smooth muscle cells (SMC) and lymphatic endothelial cells (LEC)) in iMCD-TAFRO lymph nodes. (E) For each patient per group (P1-P4) in iMCD-TAFRO (n=4) and Infl. CTRL (n=4), frequency of each stromal cell population among all non-hematopoietic/stromal cells. Volcano plot of transcription factor activity scores calculated in iMCD-TAFRO and control FRC and (G) Perivascular reticular cells (PRC).

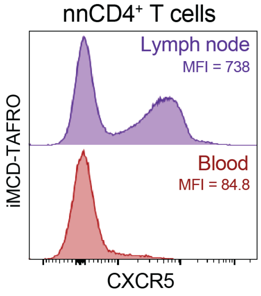


**Supplemental Figure 3. Flow cytometry histogram of CXCR5 expression.**

Flow cytometry histogram of non-naïve (nn) $CD4^+$ T cells within circulating PBMCs and from the lymph node. Median fluorescence intensity, MFI.

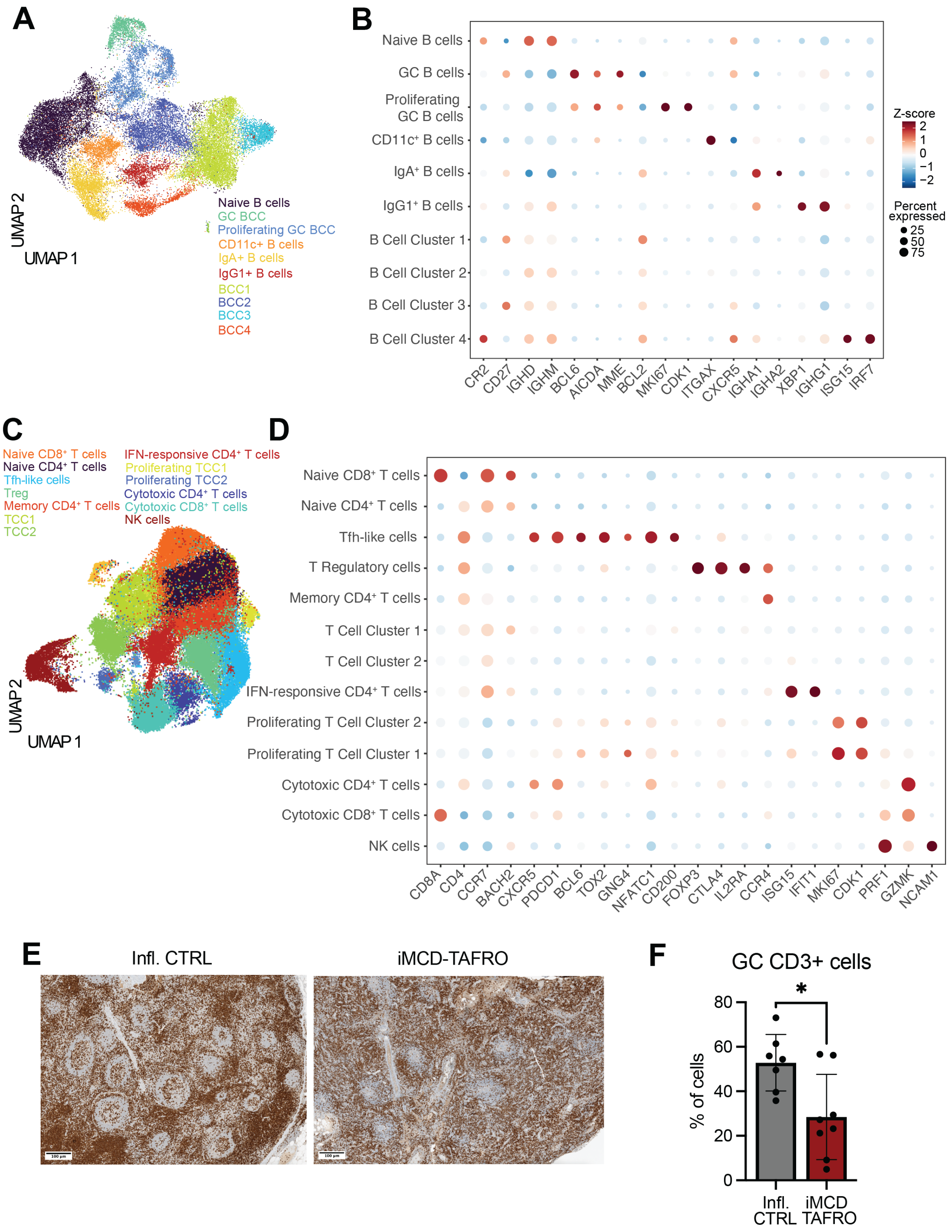


**Supplemental Figure 4. Lymphocyte characterization in iMCD lymph nodes.**

(A) UMAP of scRNAseq B cell sub-cluster. (B) Dot plot of gene expression used to identify specific B cell populations in the B cell sub-cluster with size of dot representing percent of cells expressing the gene and color representing Z score. (C) UMAP of scRNAseq T/NK cell sub-cluster and associated cell populations indicated. (D) Dot plot of specific T/NK genes that mark specific cell populations. (E) Representative images of immunohistochemistry against CD3 in inflammatory control and iMCD-TAFRO lymph nodes. (F) Quantification of immunohistochemistry of $CD3^+$ cell number in inflammatory control (n=7) and iMCD-TAFRO (n=8) lymph nodes. *$p<0.05$.

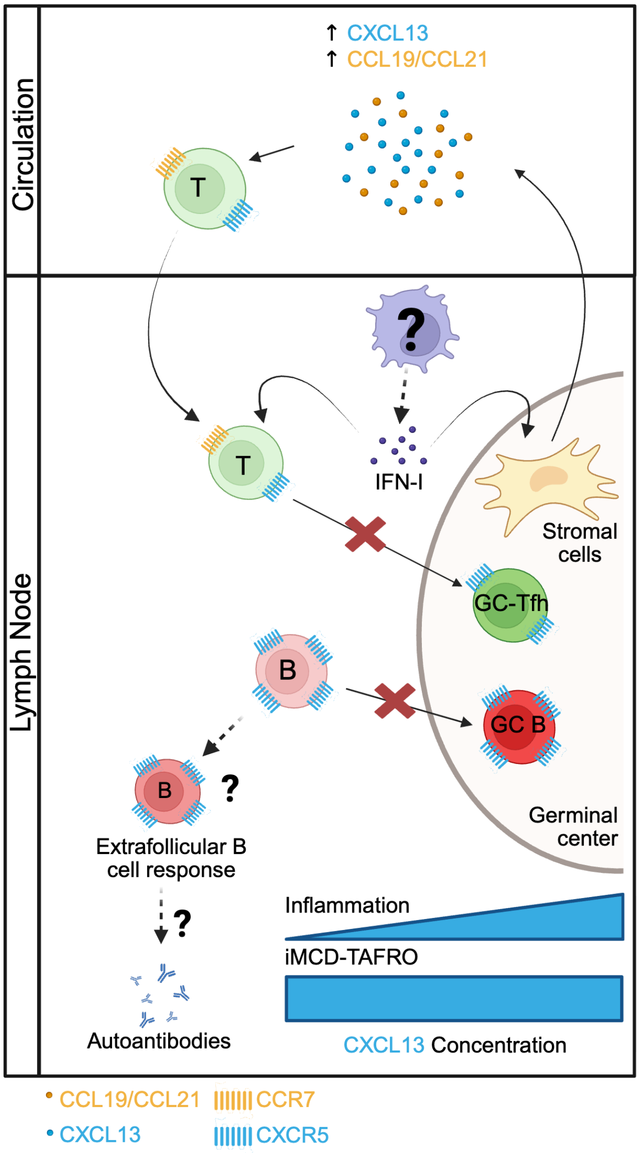


**Supplemental Figure 5. Proposed model of iMCD-TAFRO during flare.**

During active disease in iMCD-TAFRO, type I IFN in lymphoid tissue promotes chemokine production, particularly CXCL13, in lymph node stromal cells. Chemokine-responsive T and B cells follow the CXCL13-CXCR5 axis out of circulation towards heightened levels of CXCL13 in lymph node GC. After arriving in the lymph node, CXCR5-expressing T cells experience strong type I IFN signals that restrict their differentiation and movement into the GC, where CXCL13 expression typically peaks. Thus, appropriate GC responses are disrupted potentially leading to autoantibody production.
